# Synergistic interactions between *Bacteroides intestinalis* and Arabinoxylan Mitigates Inflammatory and Metabolic Dysfunction

**DOI:** 10.64898/2026.01.28.702158

**Authors:** Ziyu Zhou, Ka Lam Nguyen, Sijie Chen, Yuexi Wang, Min Li, David M. Bianchi, Weihao Ge, Shanny Hsuan Kuo, Salwa Gharieb, Emily Tung, Ronak Parmar, Jie Ji, Rohit Khorana, Ahmed Hetta, Isaac Cann, Roderick I. Mackie, Gee W. Lau, Jing Yang, Wenyan Mei

## Abstract

Insufficient dietary fiber intake is closely linked to gut microbiome dysfunction and increased risk of noncommunicable diseases. Synergistic synbiotics, pairing defined microbes with their dietary substrates, offer a precise strategy to restore microbiome function. Here, we show that pairing the human colonic commensal *Bacteroides intestinalis* with insoluble wheat arabinoxylan (inWAX) yields pronounced metabolic and anti-inflammatory benefits. Using mono-associated gnotobiotic mice and high-fat diet-induced obese mice, we demonstrate that this synbiotic enhances resistance to intestinal inflammation and improves glucose homeostasis. Mechanistically, this synbiotic increases production of 6-hydroxylated bile acids with anti-diabetic and anti-steatotic effects. It also promotes microbial transformation of phenolic compounds and redirects protein metabolism toward neuroactive metabolites. These metabolic shifts are accompanied by transcriptional remodeling in the colon, spleen and liver, involving induction of genes essential for circadian rhythm, lipid metabolism, immune defense, and bile acid 6-hydroxylation. Notably, Y chromosome-linked genes associated with epigenetic regulation and protein turnover are also induced, suggesting a potential sex-specific response to synbiotics. Together, our findings establish a mechanistic framework for targeted synbiotic interventions to combat inflammatory and metabolic disorders.

## Introduction

The global adoption of a low-fiber Western diet has fundamentally altered gut microbiota function, contributing to the escalating burden of noncommunicable metabolic and inflammatory diseases worldwide ^1^. In contrast, higher dietary fiber intake is linked to increased gut microbiome diversity and reduced risk of metabolic and inflammatory diseases ^2^. Dietary fiber has been shown to lower cholesterol, improve glucose homeostasis and weight control, and reduce the risk of colorectal cancer ^2^. Emerging evidence further implicates dietary fiber in supporting cognitive function ^3,4^. These important observations underscore the therapeutic potential of dietary fiber in preventing and managing noncommunicable diseases. Yet, the fiber-deficient Western diet has limited the daily consumption of fiber and caused the loss or reduction of many fiber-fermenting microbes in gut microbiota, thereby diminishing the benefits conferred by dietary fiber. This has spurred interest in microbiome-targeted interventions using dietary fibers (prebiotics), fiber-fermenting microbes (probiotics), and their combinations. Among these, synergistic synbiotics, which are rationally designed combinations of defined microbial strains and their preferred carbohydrate substrates, represent a particularly promising supplemental strategy. By enhancing microbial engraftment, persistence, and metabolic activity, synbiotics offer a means to restore gut microbial function and improve host metabolic outcomes in a targeted and durable manner. Despite their promise, the rational design of effective synbiotic interventions remains a significant challenge. Critical gaps remain in our understanding of microbial fiber-degrading capacities, substrate specificities, and the host-relevant bioactivities of fermentation-derived metabolites that mediate metabolic benefits.

The benefits of dietary fiber stem not only from its physicochemical properties but also from bioactive metabolites produced during its fermentation by gut commensal bacteria. Short-chain fatty acids (SCFAs) represent the most well-studied class of fiber-derived metabolites and exert broad benefits on immunity, intestinal barrier integrity, and metabolic homeostasis ^5,6^. Emerging findings, however, suggest that many health-promoting effects of dietary fiber may be attributed to microbial metabolites other than SCFAs ^7^. A better understanding of gut microbes that produce such metabolites and their associated health benefits is essential for fully harnessing the therapeutic potential of gut microbe–mediated fiber fermentation.

Here, we identify *Bacteroides intestinalis* 17393 (hereafter BI) as a key microbial mediator of dietary fiber-driven metabolic and immune benefits. BI is a member of fiber-degrading *Bacteroides* spp. of the phylum Bacteroidota, a major group of bacteria in the human adult distal gut. Originally isolated from a healthy Japanese individual^8^, this bacterium harbors esterase gene-enriched polysaccharide utilization loci (EGE-PULs) capable of sensing and degrading insoluble wheat arabinoxylan (inWAX) *in vitro* to release ferulic acid, a phenolic compound with potent anti-inflammatory and antioxidant properties ^9^. InWAX is a major plant cell wall polysaccharide abundant in cereals such as wheat, barley, and oats. It consists of a β1-4-linked xylose backbone decorated with arabinose side chains that are esterified to hydroxycinnamic acids, such as ferulic and p-coumaric acids ^10^.

Using mono-associated gnotobiotic mice and conventionally raised mice with high-fat diet–induced obesity, combined with unbiased multi-omics approaches, we demonstrate that this synbiotic confers protection against intestinal inflammation and improves glucose homeostasis. Mechanistically, it reprograms microbial metabolism to enhance the production of health-promoting metabolites, while coordinately activating transcriptional programs in the colon, spleen, and liver that reinforce immune and metabolic resilience. These findings highlight the therapeutic potential of leveraging synbiotics as a targeted strategy for the prevention and treatment of metabolic and inflammatory disorders.

## Material and Methods

### Bacteria culture

*Bacteroides intestinalis* DSM 17393 (BI) was grown anaerobically at 37°C in Brain Heart Infusion with Supplements (BHIS) medium^11^. Bacterial growth was monitored by measuring the optical density at 600 nm (OD₆₀₀). To inoculate mice, BI was cultured anaerobically until reaching log-phase growth.

### Ethics statement of mice

All mice used in these experiments were housed at the University of Illinois Urbana Champaign (UIUC) animal care facilities and cared for according to the institutional ‘Guide for the Care and Use of Laboratory Animals’. All procedures involving mouse care, euthanasia, and tissue collection were approved by the UIUC Animal Care and Use Committee (IACUC approved protocol #23096).

### Diets

BI mono-associated mice were fed either a standard chow diet (Envigo Teklad, 2019S) or custom diets containing cellulose or inWAX. Custom chow diets were formulated using an open standard diet (D11112201; Research Diets) as the base. InWAX or cellulose was added to replace the standard cellulose component, yielding diets containing 5%, 10%, or 15% inWAX (w/w). In diets containing 5% or 10% inWAX, cellulose was added proportionally to achieve a total dietary fiber content of 15% (w/w). In the diet containing 15% inWAX, cellulose was fully replaced by inWAX.

For conventionally raised mice, custom HFDs were prepared using a 60% kcal fat rodent diet (D12492; Research Diets) as the base. InWAX or cellulose was used to replace approximately 6.5% cellulose present in the standard HFD, resulting in diets containing 10% cellulose or 10% inWAX.

This diet design preserved identical macronutrient caloric composition between cellulose- and inWAX-containing diets. Detailed diet compositions are provided in Table S1. InWAX was purchased from Neogen Corporation (MI, USA; P-WAXYI; >90% purity), and cellulose was supplied by Research Diets, Inc.

### BI mono-association in mice fed standard chow diet

GF mice fed a standard chow diet were orally gavaged every other day with 200 µL of BHIS medium containing live BI (2.24 × 10⁸ CFU/mL), heat-killed BI, or BHIS medium alone (vehicle control). The absence of viable bacteria in heat-killed BI preparations was confirmed by anaerobic incubation of BI preparations in BHIS broth at 37°C, with growth monitored for up to 3 days. Viability of BI in mice inoculated with live BI was confirmed midway through the treatment period by quantifying colony-forming units (CFUs) from fecal samples using conventional plating on BHIS agar.

The identity and specificity of BI in colonized mice were validated by sequencing PCR amplicons generated from fecal microbial DNA using both universal 16S rRNA primers and BI strain–specific primers (sequences are provided in Table S2). Following euthanasia, intestinal length and cecum weight and length were measured. Intestinal tissues were either flash-frozen, fixed in 4% paraformaldehyde for histological analyses, or fixed in Methacarn solution (60% methanol, 30% chloroform, 10% glacial acetic acid) for mucus architecture assessment. Goblet cells and mucus were visualized in Methacarn-fixed, paraffin-embedded colon sections using Alcian Blue staining (pH 2.5)^12^ or Alcian Blue pH 2.5 - Brown Hopps Gram staining with slight modification^13^. Specifically, 95% ethanol was used in place of Cellosolve as the differentiating solution.

### Goblet cell quantification

A defined region of mouse middle colon was selected for analysis. The relative number of goblet cells per crypt was determined by calculating the ratio of goblet cells to the total number of colonic epithelial cells within each crypt. At least 30 measurements per mouse were collected.

### Colon mucus layer quantification

Measurements were performed using ImageJ in distal colon regions exhibiting a continuous mucus layer ≥100 µm in length, with an intact epithelial lining and clearly defined luminal content stratification. Mucus thickness was quantified as the perpendicular distance spanning the visibly firm, dense mucus layer. For each qualifying region, a minimum of three perpendicular measurements were obtained at positions spaced 50 µm apart along the length of the mucus layer. Serial sections were collected at 50 µm intervals from each mouse to ensure that at least 30 measurements were obtained from comparable anatomical regions at different depths within the same animal. Measurements were averaged in sets of six, and these means were pooled for statistical comparisons between experimental groups.

### BI mono-association in mice fed chow diet containing inWAX or cellulose

GF mice were acclimated to diets containing inWAX or cellulose for 5 days prior to BI inoculation. Mice were then orally gavaged with BI (200 µl at the concentration of 2.24×10⁸ CFU/mL) every day. Body weight was monitored daily throughout the experiment. The viability and specificity of BI in colonized mice were validated by conventional plating and PCR-based sequencing as described above. Following euthanasia, tissues and serum were collected. Serum samples were processed for ferulic acid measurement or untargeted metabolomic profiling. Intestines were either flash-frozen or fixed in 4% paraformaldehyde for histological evaluation.

### BI inoculation in mice fed HFD diet with or without inWAX or cellulose

14-week-old conventionally raised C57BL/6J mice preconditioned with an HFD for 8 weeks were purchased from The Jackson Laboratory (strain no. 380050). Mice were acclimated to HFDs with or without inWAX or cellulose for 5 days prior to BI inoculation. Mice were then orally gavaged with BI every other day. BI abundance and specificity in colonized mice were confirmed by PCR and PCR-based sequencing. Glucose tolerance test was measured at the end of feeding trial as described below. At the experimental endpoint, mice were fasted overnight and subsequently refed with their respective diets. Fresh fecal samples were collected 2 hours after refeeding for targeted bile acid quantification and metabolomic analyses. Following euthanasia, tissues were collected for downstream analyses.

### Glucose tolerance test

Mice treated with BI and fed HFDs, with or without cellulose and inWAX, were fasted overnight and administered glucose (1 g/kg body weight) via intraperitoneal injection. Blood glucose levels were measured from tail vein blood at baseline (0 minute) and at 15-, 30-, 60-, 90-, and 120-miniutes post-injection.

### DSS-induced colitis

BI mono-associated mice received either 15% inWAX or 15% cellulose were treated with 2% (w/v) dextran sulfate sodium (DSS) in drinking water for 5 days (referred to as inWAX;DSS+ and cellu;DSS+, respectively). The control mice received 15% inWAX without DSS treatment (inWAX;DSS-). All mice were maintained on their respective diets throughout the DSS or water treatment period. Body weight was monitored daily as clinical readouts of colitis. On day 5 of DSS treatment, mice were euthanized for tissue collection. A range of physical parameters were evaluated, including colon length, spleen weight, and spleen length.

For histological analyses, tissues were fixed in 4% paraformaldehyde and paraffin processed for histomorphology. Unless otherwise specified, tissue sections were cut at 5 μm thickness. Tissue sections were stained with hematoxylin and eosin (H&E) for morphology or Masson’s Trichrome for fibrosis ^12^. Images were acquired using a Keyence BZ-X810 microscope. Histopathological scores were blindly assessed as described previously.^14^ Briefly, each colon was assessed for four parameters: inflammatory cell infiltration, epithelial architecture destruction, crypt dilatation, and fibrosis within the mucosa, submucosa, and serosa. Significant histopathological features, such as loss of crypt structure, goblet cell depletion, mucosal erosion, and submucosal fibrosis were recorded to further distinguish the severity of colonic injury and inflammation.

### Immunohistochemistry

For Immunohistochemistry, paraffin sections were deparaffinized and subjected to heat-induced antigen retrieval using sodium citrate buffer (10 mM sodium citrate, 0.05% Tween-20, pH 6.0) for 20 minutes. Sections were incubated overnight at 4°C with a rabbit anti-MUC2 antibody (Genetex, GTX100664) diluted in PBS containing 5% normal serum and 1% BSA. Detection was performed using Alexa Fluor 488-conjugated goat anti-rabbit secondary antibody (Thermo Fisher Scientific, A32731TR). Nuclei were counterstained with 4’,6-diamidino-2-phenylindole (DAPI) and imaged using a Nikon A1R confocal microscope.

### Fluorescence in-situ hybridization (FISH)

A BI-specific probe was designed according to established guidelines^15^. Following the reported nomenclature^16^, the probe was designated S-S-B.int-0461-a-S-22 and is referred to herein as “Bint461” for clarity and convenience. The probe was 5’-labeled with sulfonated fluorescein 488 (Sigma-Aldrich). To assess specificity, FISH was performed on pure cultures of BI and *Escherichia coli* as described ^17^. Localization of BI in colon sections from mono-associated mice was assessed using the Bint461 probe. Total bacterial presence was simultaneously detected using a universal bacterial FISH probe, S-D-Bact-0338-a-A-18 (Eub338), which was 5’-labeled with Texas Red as a positive control^18^. The NON-EUB338 probe, designed as the reverse complement of EUB338, was included as a negative control. Slides were counterstained with DAPI and images were acquired using a Nikon A1R confocal microscope. The images are shown in Figure S1. Probe sequences are provided in Table S2.

### Ferulic acid measurements

Absolute quantification of ferulic acid was done using Liquid Chromatography-tandem Mass Spectrometry (LC-MS/MS) by the Carver Metabolomics Core Facility at UIUC. Briefly, serum samples were spiked with a mixture of deuterium- and stable isotope-labeled internal standards prior to alkaline hydrolysis with NaOH, which facilitated the release of ferulic acid from its conjugated forms. Total ferulic acid content, comprising both free ferulic acid and that liberated from conjugates, was subsequently quantified.

### Untargeted metabolomics assay

Samples were spiked with isotope-labeled internal standards prior to extraction and processed for untargeted metabolomics by the Carver Metabolomics Core at the Roy J. Carver Biotechnology Center at UIUC. Fecal samples were normalized to input dry fecal weight. Metabolomic profiling was performed using a Dionex Ultimate 3000 UHPLC system coupled to a Q-Exactive mass spectrometer. Data were acquired in both positive and negative electrospray ionization modes. Raw LC–MS data were processed using MS-DIAL (v4.90) for feature detection, alignment, adduct annotation, and compound identification based on accurate mass, MS/MS spectra, and retention time matching to an in-house standards library and public databases (MoNA, MassBank, GNPS, METLIN, and NIST20). Features with a sample-to-blank ratio <10 were excluded, and metabolites were assigned confidence levels according to Metabolomics Standards Initiative (MSI) guidelines. Positive and negative ionization datasets were merged, duplicate features were removed using quality control metrics, and peak intensities were normalized to internal standards. Principal component analysis was used for data quality assessment and outlier detection.

### Metabolomics Data Processing and Statistical Analysis

Pairwise comparisons between experimental groups were performed using a custom function that calculated log2 fold changes and p-values for each metabolite. Zero values were excluded prior to analysis. Log2-transformed metabolite intensities were compared between groups using t-tests assuming unequal variance. Significant metabolites were defined based on both p-value (<0.05) and log2 fold change thresholds. Volcano plots were generated to highlight significant up- and down-regulated metabolites. Heatmaps of selected metabolites were created with row-standardization (Z-score) applied.

To investigate shared and unique metabolites across group comparisons, Venn diagrams and UpSet plots were constructed. Metabolites were classified as “unique” or “shared” based on their occurrence across comparisons, and compound class information was retrieved from external databases. A Pie chart was generated to visualize the distribution of metabolite categories for uniquely upregulated metabolites in inWAX-treated mice. All data processing, statistical analyses, and visualizations were performed in R (v4.5.1) using the following packages: ggplot2 (v4.0.1)^19^, ggrepel (v0.9.6)^20^, ComplexHeatmap (v2.16.0)^21^, patchwork (v1.3.2)^22^, dplyr (v1.1.4), tidyr (v1.3.1), tibble (v3.3.0), and readxl (v1.4.5) ^19^.

### Targeted bile acid assay

Samples were analyzed using LC-MS by the Carver Metabolomics Core of the Roy J. Carver Biotechnology Center, UIUC. The bile acid standard solution (Cayman chemical, Ann Arbor, Michigan, USA), containing 1 mg/mL of each of targeted bile acids was used to create a calibration curve for quantification. At the beginning of the extraction process, surrogate internal standards were spiked into each sample. Instrument internal standards (CUDA, PUHA, 0.1 µg/mL) were included in the reconstitution solution. Chromatography was performed on an Agilent 1290 Infinity II UHPLC system, with Waters Acquity C18 BEH 2.1 x 100mm, 1.7um; flow rate 450 μL/min. Data was collected on an Agilent 6495C triple quadrupole mass spectrometer in negative MRM mode. Peak integration and quantitation using calibration curves adjusted for internal standards were done using Mass Hunter Quant 12.1. Significantly altered bile acids were identified using a two-tailed Mann–Whitney U test. A p-value < 0.05 was considered statistically significant.

### RNA-sequencing analysis

RNAs were extracted from the colons and spleens using PureLink RNA Mini Kit (Thermo Fisher Scientific, 12183025). RNA-seq libraries were constructed and sequenced at the UIUC Biotechnology Center High-Throughput Sequencing Core. Each library generated over 80 million single-end reads. Adaptor sequences and low-quality bases were removed from the raw sequencing data using Trimmomatic v0.39.^23^ Transcript abundance was quantified using Kallisto v0.44.0 ^24^, aligning reads to the mouse transcriptome (GRCm39, Ensembl). Data quality and sample consistency were evaluated through principal component analysis (PCA) and hierarchical clustering based on transcript quantifications. Transcript-level expression values corresponding to the same gene were then aggregated to obtain gene-level expression estimates. Differentially expressed genes (DEGs) were identified using DESeq2 v2_1.46.0 ^25^, with a significance threshold of adjusted p-value < 0.05 and |log2 fold change| > 1. Gene Ontology (GO) functional enrichment analysis was performed using Metascape identify significantly enriched biological processes, molecular functions, and cellular components^26^.

### Immune Cell Fraction Analysis

Immune cell composition was assessed from bulk RNA-seq data using a deconvolution-based approach implemented in the R package immunedeconv^27^. Mouse gene symbols were converted to their human orthologs using the convert-human-mouse-genes function. Immune cell fractions were inferred using quanTIseq ^28^, a constrained least squares regression method, with all expressed genes (TPM as the unit) to deconvolute the relative proportions of immune cell populations within each sample.

### Quantitative Real-Time PCR (qPCR)

RNAs were extracted using TRIzol reagent according to standard protocols or using PureLink RNA Mini Kit. Real-time PCR reactions were performed blindly in duplicate using the SYBR green master mix. Primer sequences are shown in Table S2.

### Statistical analysis

Graphs were generated using GraphPad Prism software or R. Unless otherwise noted, statistical significance was determined as follows: differences among three groups were analyzed using one-way ANOVA followed by Tukey’s multiple comparisons test or Šidák post-hoc test, or by Kruskal-Wallis test followed by Dunn’s test. Comparisons of gene expression between two groups were performed using a two-tailed unpaired *t* test. Differences in serum ferulic acid levels between selected two groups were assessed using a two-tailed Mann–Whitney U test.

## Results

### BI promoted colon mucus layer formation

To determine the physiological impact of BI, we inoculated viable BI and heat-killed BI to germ-free (GF) mice fed a standard chow diet (Figure 1A, Materials and Methods). The GF control mice were inoculated with vehicle alone (culture medium). Comparison of physiological responses to viable versus heat-killed BI allowed us to distinguish effects mediated by BI-derived metabolites, as heat-killed bacteria are incapable of metabolite production. Viable BI significantly reduced the ratios of cecum size and weight to the body weight when compared to the vehicle control and heat-killed BI but had no effect on body weight and intestine lengths (Figure 1B and S2A). Both viable BI and heat-killed BI significantly increased the percentages of goblet cells in the colon, duodenum, and ileum but not in the jejunum when compared to the vehicle control (Figure 1C and S2B). However, only mice inoculated with viable BI developed a substantially thicker mucus layer in the colon (Figure 1D and S2C). These findings suggest that both BI cell wall components and metabolites jointly promote region-specific goblet cell differentiation, whereas mucus layer formation is stimulated by BI-derived metabolites.

**Figure 1.**
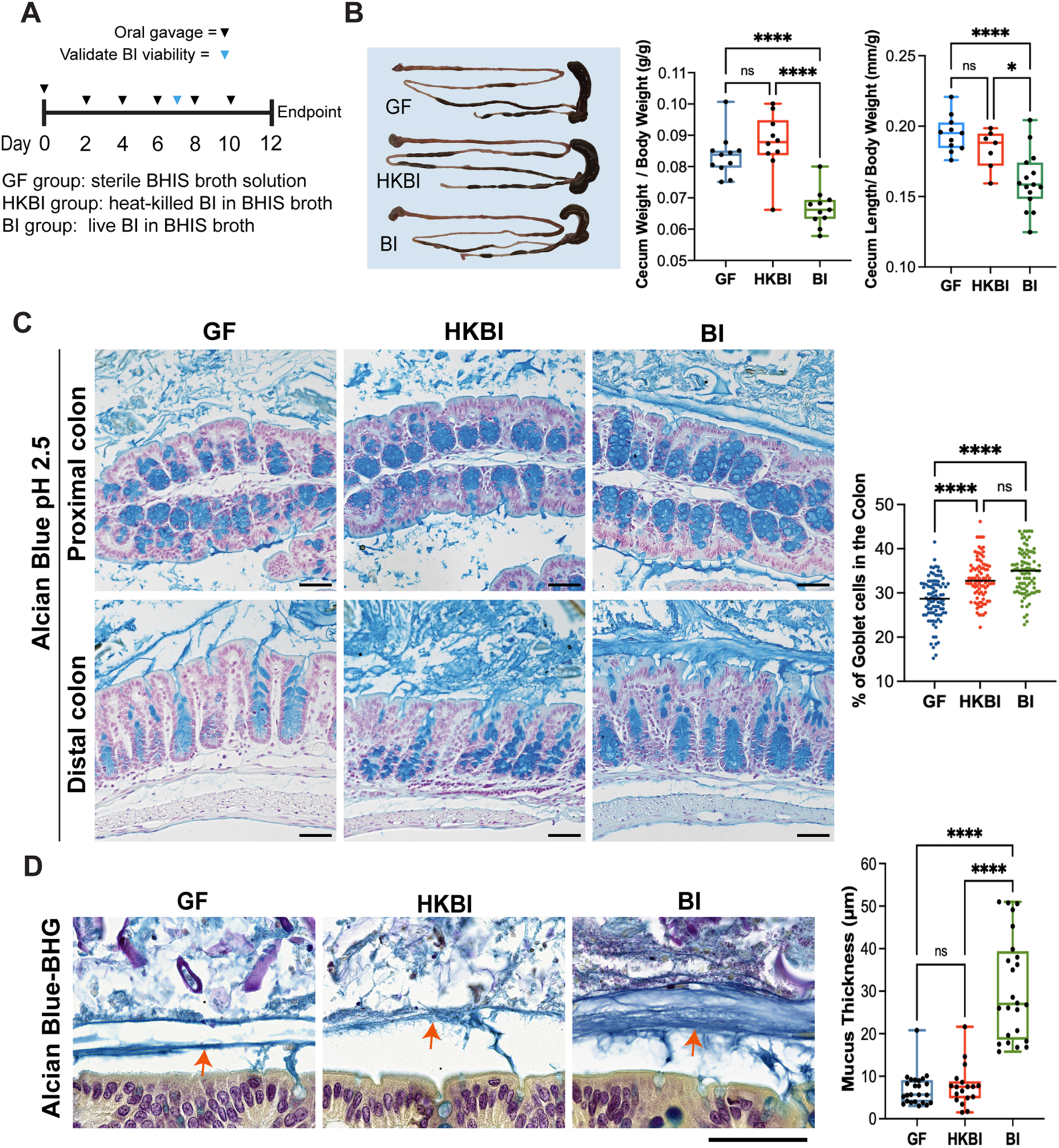
BI mono-association reduces mouse cecum size and enhances intestinal mucus secretion. (A) Schematic of the experimental design and treatments for each mouse group. (B) Representative images of the small intestine, cecum, and colon from mice in each group, with the ratio of cecum size and weight to body weight quantification shown on the right. (C) Alcian blue staining shows goblet cells in the proximal and distal colons of BI-inoculated mice. Quantification of percentages of goblet cell is shown on right. (D) Alcian blue-BHG staining showing increased thickness of colon mucus layer (red arrows) in BI-inoculated mice relative to germ-free (GF) mice and heat-killed (HK)-BI-inoculated mice. quantification is shown on the right. Scale bars, 50 μm. **** p < 0.0001; ns, not significant.

### BI-mediated fermentation of inWAX increased systemic level of ferulic acid

We next examined BI’s capability and capacity of fermenting inWAX and releasing ferulic acid *in vivo*. GF mice were inoculated with viable BI or vehicle and fed a custom diet containing 5%, 10%, or 15% inWAX, or a control diet containing the same ingredients except that inWAX was replaced with 15% cellulose (Figure 2A). In the diets containing 5% or 10% inWAX, cellulose was proportionally added to make the total fiber 15% (Table S1). Cellulose was used as a control for non-specific fiber effect because BI lacks the enzymatic capacity to degrade it^9^. It also served as a control for slightly reduced calorie density because of the introduced fiber. Systemic ferulic acid levels in mouse serum were compared across groups. As shown in Figure 2B, all three concentrations of inWAX in the diet led to a significant increase in systemic levels of ferulic acid when compared to cellulose, and the level of circulating ferulic acid correlated with the amount of inWAX intake. However, when the inWAX concentration was increased from 10% to 15%, there was no further significant increase in ferulic acid levels, suggesting that 10% inWAX may represent a dose close to the saturation point of BI’s fermentation capacity. These results demonstrate that BI can ferment inWAX *in vivo* to release ferulic acid and elevate its systemic level, highlighting its anti-inflammatory and antioxidant potential.

**Figure 2.**
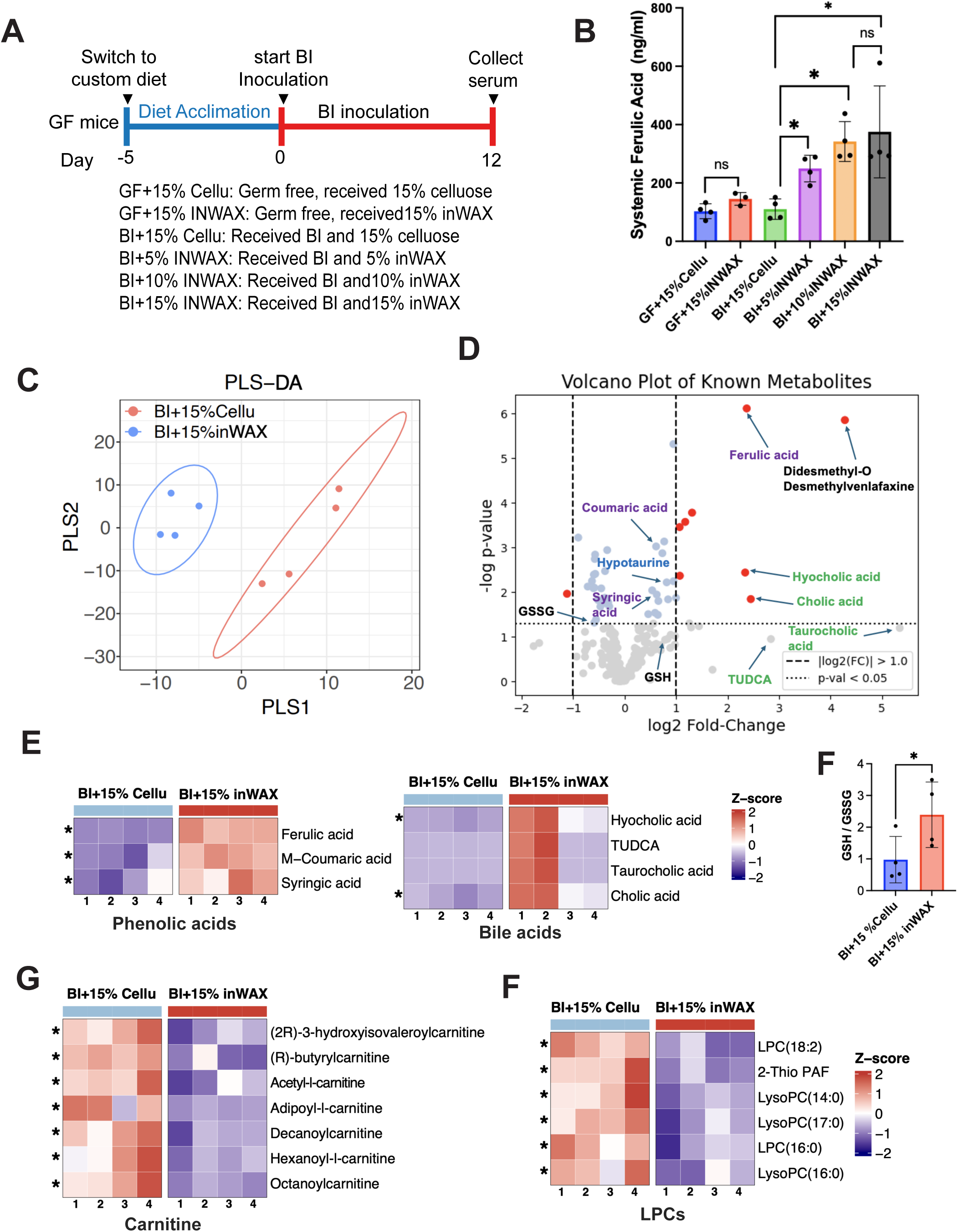
BI and inWAX increased beneficial bile acids and phenolic compounds. (A) Schematic of the experimental workflow and treatments for each mouse group. (B) Comparison of systemic levels of ferulic acid across 6 groups of mice. (C) Partial least squares discriminant analysis (PLS-DA) showing distinct circulating metabolite profiles induced by inWAX versus cellulose in BI mono-associated mice. (D) Volcano plot showing differentially abundant circulating metabolites in inWAX-fed mice relative to cellulose-fed mice; bile acids are highlighted in green and phenolic compounds in purple. (E) Heatmaps showing inWAX-induced increases in phenolic compounds and bile acids. (F) Comparison of the GSH/GSSG ratio. * p < 0.05, one-tailed T-test. (G) Heatmaps showing inWAX-induced reductions in lysophosphatidylcholines (LPCs) and carnitine derivatives. An asterisk (*) on the heatmaps indicates statistical significance.

### BI-mediated fermentation of inWAX generated disease-protecting blood metabolome

To unbiasedly evaluate the impact of BI-mediated fermentation of inWAX on host metabolism, we performed an untargeted metabolomics analysis using serum samples from BI-inoculated mice fed 15% inWAX or 15% cellulose. Mice receiving inWAX diet showed a very distinct circulating metabolic profile from those receiving cellulose (Figure 2C), indicating that metabolites released from BI-mediated fermentation reshaped host blood metabolome. 22 known metabolites were significantly elevated by inWAX (Figure 2D, Table S3). Among them, ferulic acid is one of the most elevated metabolites, which is consistent with the observation from the ferulic acid quantification assessment (Figure 2B). In addition to ferulic acid, two additional phenolic acids that have antioxidant and anti-inflammatory properties, m-coumaric acid and syringic acid, are also significantly increased (Figure 2D and 2E). This is accompanied by an elevated ratio of reduced glutathione (GSH) to oxidized glutathione (GSSG) (Figure 2F), a hallmark of reduced oxidative stress and an increased antioxidant capacity.

Notably, four of the six most highly elevated metabolites in inWAX-fed mice were bile acids, including cholic acid (CA), taurocholic acid (TCA), tauroursodeoxycholic acid (TUDCA), and hyocholic acid (HCA) (Figure 2D and 2E). The increases in TCA and TUDCA narrowly missed statistical significance due to substantial inter-individual variability within the inWAX-treated group. Importantly, both TUDCA and HCA have well-established anti-diabetic properties^29–31^. TUDCA is clinically used for treating liver cholangitis and has recently been shown to reduce intestinal inflammation in both mouse models and patients with IBD ^32^ ^33^.

The metabolite that showed the highest increase in inWAX-fed mice is D,L-N,N-didesmethyl-o-desmethylvenlafaxine, a derivative of venlafaxine. Venlafaxine is a serotonin-norepinephrine reuptake inhibitor and has been clinically used for treating depression and anxiety ^34^. The elevation of this compound suggests the presence of an as-yet-uncharacterized venlafaxine-like metabolite induced by BI-mediated inWAX fermentation.

23 known metabolites are significantly decreased in inWAX-treated mice (Table S3). Intriguingly, more than 56% of them are involved in lipid metabolism, including 7 derivatives of carnitine and 6 members of lysophosphatidylcholines (LPCs) (Figure 2G). Derivatives of carnitine act as a transport system to move activated fatty acids into mitochondria for β-oxidation, which is a primary energy-generating process. LPCs are bioactive signaling molecules derived from the partial hydrolysis of phosphatidylcholines and are involved in lipid signaling and cell membrane remodeling. The decrease in the circulating levels of these metabolites is presumably an indicative of improved lipid metabolism and energy homeostasis through a gut-liver-metabolism axis.

Together, these findings demonstrate that supplementation with BI and inWAX elevates systemic levels of phenolic compounds and bile acids with anti-inflammatory, antioxidant, and anti-diabetic activities, improves cellular redox balance, and reshapes lipid metabolism.

### BI-mediated inWAX fermentation shifted the colonic transcriptome

Given that the colon is directly exposed to microbial metabolites, we next assessed the effect of BI-mediated inWAX fermentation on the colon transcriptome. 15% inWAX-treated mice exhibited 17 upregulated genes in the colon when compared to 15% cellulose-fed mice (Figure 3A, Table S4). Strikingly, the gene that is mostly upregulated is *cyclic AMP responsive element–binding protein 3–like 3*(*Creb3l3*), a transcriptional factor that promotes triglyceride metabolism. Loss-of-function *Creb3l3* variants in humans have been associated with hypertriglyceridemia, whereas overexpression of *Creb3l3* reduces arteriosclerotic lesions in mouse models of arteriosclerosis ^35–38^. Thus, an elevation of *Creb3l3* underscores the potential of BI and inWAX in protecting against hypertriglyceridemia and arteriosclerosis.

**Figure 3.**
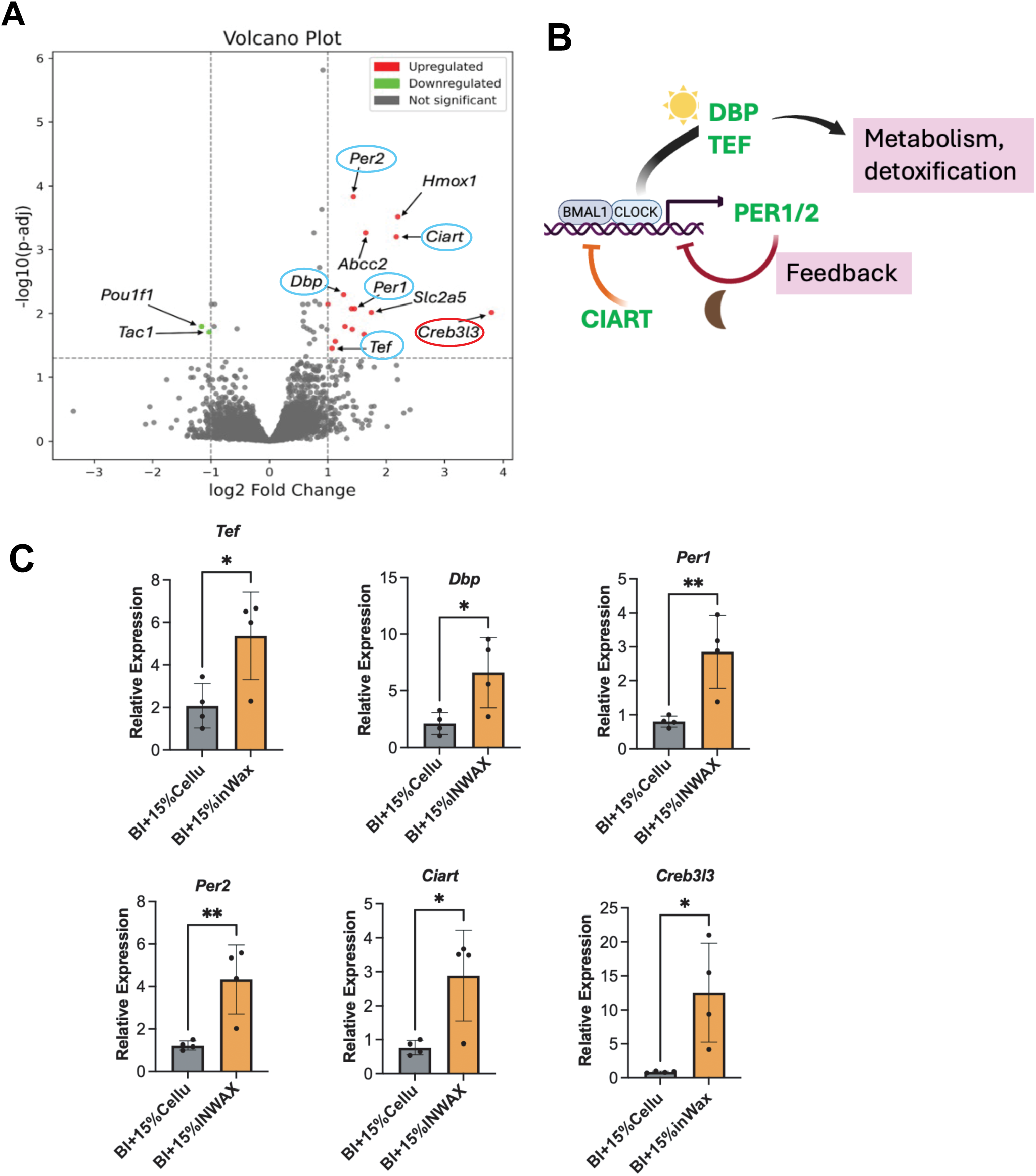
BI and inWAX activates genes essential for circadian regulation and triglyceride metabolism in the colon. (A) The volcano plot showing differentially expressed gene in the colon of inWAX-fed mice relative to the cellulose-fed mice. Circadian regulators are circled in blue and *Creb3l3* in red. (B) Schematic illustrating the roles of upregulated circadian regulatory genes within the circadian clock network. (C) Quantitative RT-PCR validation of selected upregulated genes. Expression levels were normalized to *Gapdh*. Each symbol represents an individual mouse and bars indicate mean values. * p < 0.05; ** p < 0.01.

Surprisingly, 5 increased genes are core regulators of circadian rhythms (Figure 3A). *Per1*, *Per2,* and *Ciart* (also called *Chrono*) are key components of a negative feedback loop in the circadian control by inhibiting the activity of the CLOCK–BMAL1 complex, the master transcriptional activator of clock genes, whereas *Dbp and Tef* are circadian clock-controlled transcription factors in the PAR bZip family that mediate circadian clock output pathways (Figure 3B) ^39–41^. Using quantitative real-time PCR, we validated the increased expression of these genes in the colons of inWAX-treated mice (Figure 3C).

Other upregulated genes include *Hmox1*, *Fkbp5,* and *Hif3a* that respond to stress and hypoxia ^42–44^, *Slc2a5* and *Abcc2* for membrane transport^45,46^, *Kyat1* for the tryptophan Kynurenine Pathway, and *Tsku* and *Lefty1* for tissue development and homeostasis ^47,48^.

Only two genes were downregulated in inWAX-treated mice, including *Pou1f1*, a transcriptional factor with unknown intestinal function, and *Tac1* (Figure 3A), which encodes the precursor for neuropeptides tachykinins that regulate intestinal motility, secretion, and vascular functions ^49^.

Together, these findings indicate that BI-mediated fermentation of inWAX influences intestinal function by activating genes implicated in the triglyceride metabolism and intestinal circadian clock.

### BI-mediated inWAX fermentation protects against colitis

Given that BI-mediated fermentation of inWAX increased systemic levels of anti-inflammatory and antioxidant metabolites, we investigated its protective effect against dextran sulfate sodium (DSS)-induced colitis, a major type of IBDs. BI mono-associated male mice received 15% inWAX or 15% cellulose were challenged with 2% DSS to induce colitis (Figure 4A, thereafter inWAX:DSS+ and Cellu;DSS+, respectively). Male mice treated with 15% inWAX without DSS served as controls (inWAX;DSS-). InWAX:DSS+ mice exhibited significantly less body weight loss in response to DSS treatment when compared to Cellu;DSS+ mice (Figure 4B). At the end of the treatment, Cellu:DSS+ mice showed splenomegaly because of inflammation. This splenomegaly was substantially attenuated in inWAX;DSS+ mice (Figure 4C), suggesting reduced systemic inflammation. Consistently, inWAX:DSS+ mice displayed significantly lower colonic histopathological scores, characterized by reduced crypt dilation, fibrosis, and inflammatory infiltrates (Figure 4D).

**Figure 4.**
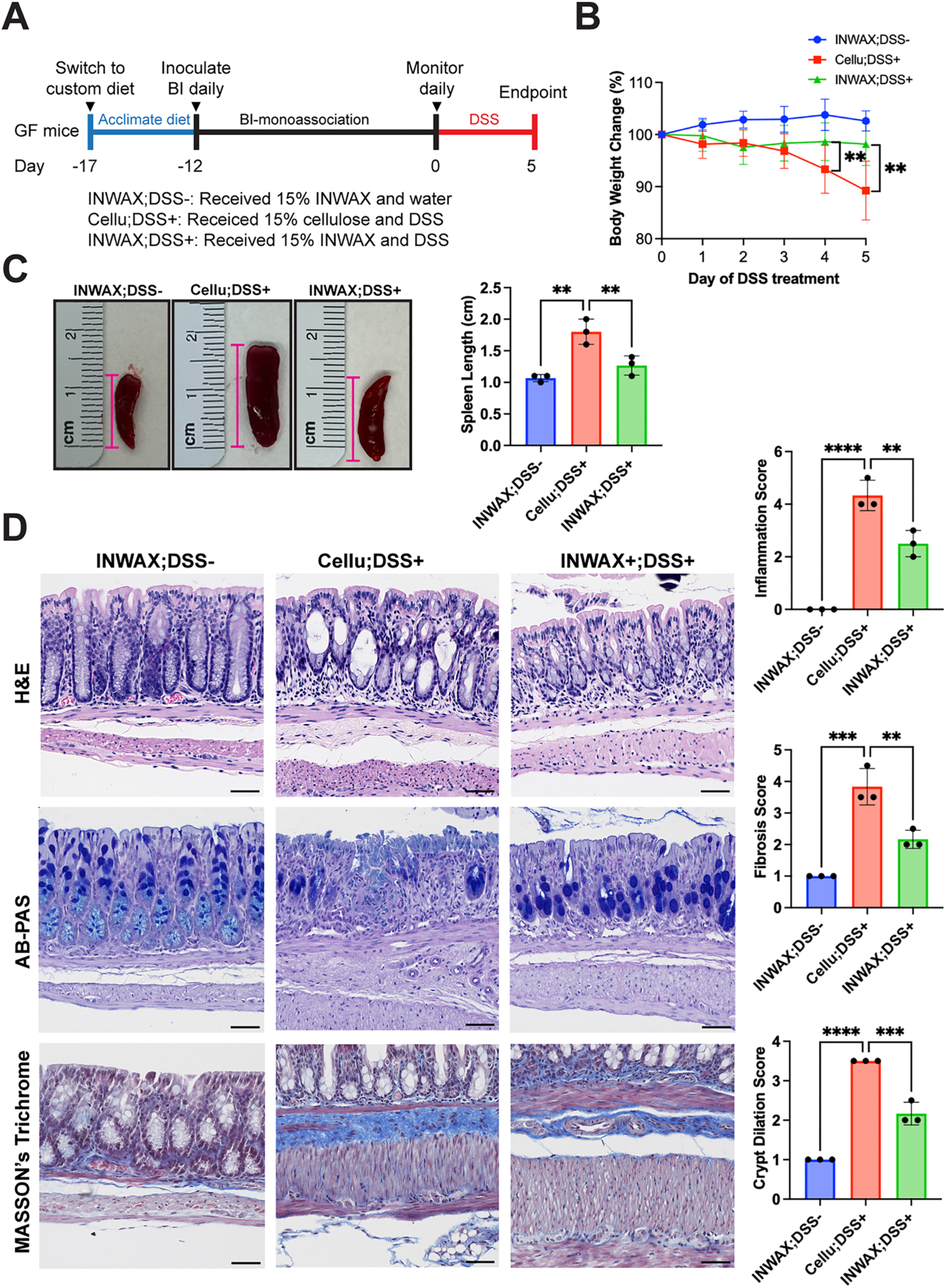
BI and inWAX ameliorates DSS-induced colitis. (A) Experimental schematic showing dietary acclimation, BI inoculation, and DSS treatment. Germ-free mice were acclimated to customized diets for 5 days, inoculated with BI, and treated with or without DSS for an additional 5 days. Compared with the cellu;DSS+ group, mice in the inWAX;DSS+ group exhibited reduced body weight loss (B), decreased splenomegaly (C), and lower histopathological scores for inflammation, crypt dilation, and fibrosis (D). Masson’s Trichrome staining (blue in D) marks fibrosis. Bars represent 50 µm.

Collectively, these results strongly indicate that BI-mediated fermentation of inWAX confers protection against DSS-induced colitis.

### inWAX fermentation induced protective colon transcriptome against DSS

To elucidate the molecular basis of colitis resistance conferred by BI and inWAX, we analyzed DSS-induced transcriptional changes in the colon. Relative to Cellu;DSS+ mice, inWAX;DSS+ mice exhibited upregulation of 52 genes (Figure 5A, Table S4). This includes 7 genes involved in innate immune defense (*Reg3b*, *Retn*, *Apol7c*, *Cd177*, *Clec14a*, *Clec2e*, *Mafb*). *Reg3b* was among the most strongly induced genes, and it encodes an antimicrobial lectin that limits bacterial translocation and protects against DSS-induced colitis ^50–52^, suggesting enhanced host defense to DSS-induced colitis.

**Figure 5.**
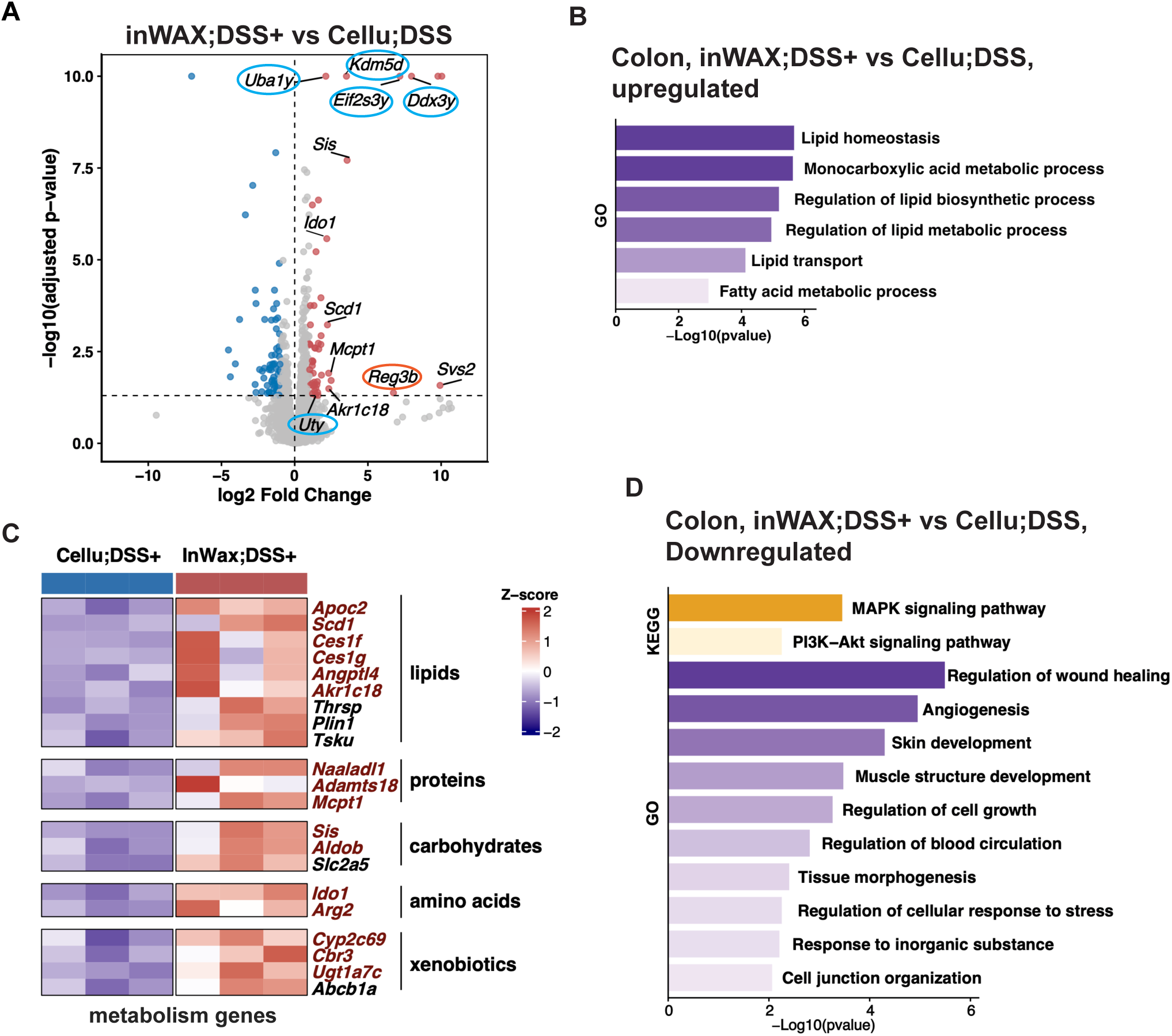
BI and inWAX remodels the colonic transcriptome in response to DSS. (A) Volcano plot showing differentially expressed genes in the colons of inWAX-fed mice relative to cellulose-fed mice following DSS treatment. Y chromosome-linked genes are circled in blue and *Reg3b* is circled in red. (B) Gene Ontology (GO) analysis showing enrichment of lipid metabolism pathways among upregulated genes. (C) Heatmap showing upregulated metabolic genes. (D) GO analysis showing enrichment of wound-healing pathways among downregulated genes.

Surprisingly, inWAX treatment induced a set of genes classically associated with male fertility, including *Svs2*, *Ddx3y*, *Uty* (*Kdm6c*), *Kdm5d*, *Eif2s3y*, and *Uba1y* (Figure 5A). Five of these genes are Y chromosome-linked and function in RNA translation (*Eif2s3y*, *Ddx3y*), epigenetic regulation (*Uty*, *Kdm5d*), and protein ubiquitination (*Uba1y*) ^53–57^. *Svs2* is not Y-linked and encodes a secreted seminal vesicle protein that protects sperm from immune attack^58^, with an unknown role in the intestine. This interesting finding suggests that BI-mediated inWAX fermentation induces sex-based immune modulation in the colon in response to DSS insult.

Notably, nearly half of the upregulated genes encoded metabolic regulators involved in lipid, protein, carbohydrate, and xenobiotic metabolism (Figure 5B and 5C). Most of them encode enzymes or regulators of enzymes, underscoring a broad metabolic reprogramming of the colon. Among them, lipid metabolic pathways show the strongest enrichment (Figure 5B and Table S5), spanning multiple layers of lipid metabolism regulation, including lipid synthesis and desaturation (*Thrsp*, *Scd1*) ^59,60^, storage (*Plin1*) ^61^, hydrolysis and mobilization (*Apoc2*, *Angptl4*, *Ces1f*, *Ces1g*) ^62–64^, as well as extracellular regulation of lipid metabolic signaling (*Tsku*). ^65^ This finding highlighted a coordinated lipid metabolic reprogramming in the colon.

Importantly, many of these metabolism regulatory genes possess immune-regulatory functions. For instance, *Scd1* encodes Stearoyl-CoA desaturase-1 that converts saturated fatty acids into monounsaturated fatty acids ^60^, and can attenuate DSS-induced colitis ^66^. *Ido1* modulates immune responses by catabolizing tryptophan^67–69^. *Mcpt1* encodes a Mast cell protease that is predominantly expressed in intestinal mucosal mast cells and has multifaced roles in inflammation ^70^. *Arg2* encodes a mitochondrial enzyme involved in arginine metabolism and nitric oxide synthesis, and can influence immune cell function and suppress inflammation ^71–74^. *Vnn1* encodes a pantetheinase that converts pantetheine to pantothenate and cysteamine to enhance SCFA production and protect against DSS-induced colitis^75^. *Angptl4* represses lipoprotein lipase and is also immune regulatory ^63^.

Consistent with attenuated inflammatory stress in inWAX-treated mice, we observed significant downregulation of genes associated with wound healing, stress responses, and cell junction assembly, along with reduced activities in MAPK signaling and PI3K-AKT signaling (Figure 5D, Table S5).

Collectively, these data indicate that BI-mediated fermentation of inWAX induces a coordinated transcriptional program that couples metabolic reprogramming with enhanced mucosal immune protection. This finding reveals a central role for gut microbiota-driven metabolic remodeling in promoting intestinal resiliency to inflammatory injury.

### inWAX fermentation induced protective spleen architecture and transcriptome against DSS

The spleen is a major secondary lymphoid organ that integrates hematopoietic and immunological functions. Given our observation that BI-mediated fermentation of inWAX markedly attenuated DSS-induced splenomegaly, we next examined the impact of BI-mediated inWAX fermentation on splenic architecture and transcriptional programs. Compared with cellulose-treated controls, inWAX-treated mice exhibited a pronounced expansion of the white pulp (Figure 6A), the lymphocyte-enriched compartment of the spleen, indicative of enhanced adaptive immune cell abundance. This was confirmed by transcriptome profiling, which showed broad activation of genes essential for B and T cell proliferation and differentiation, activation, immunoglobulin production, and cytokine signaling, including those encoding components of Th1, Th2, Th17, and JAK-STAT signaling pathways (Figure 6B, Table S4 and S5). In contrast, genes suppressed by inWAX treatment were enriched for DNA replication and cell cycle pathways (Figure 6C, Table S4 and S5), which presumably attenuated splenomegaly.

**Figure 6.**
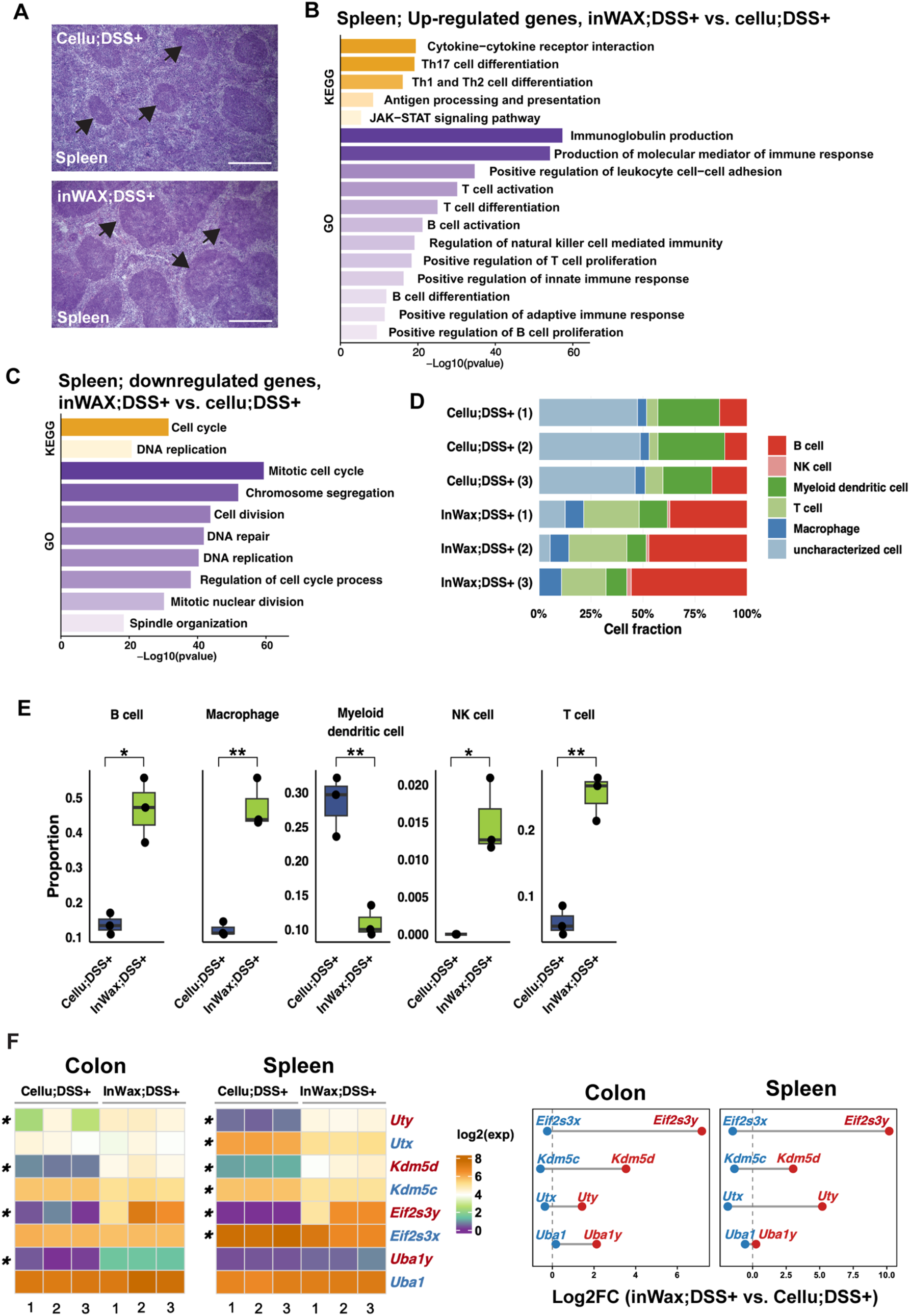
BI and inWAX reshape the spleen immune cell landscape in response to DSS. (A) H&E-stained spleen sections showing enlarged white pulp regions in inWAX-treated mice (arrows). (B) GO analysis showing enrichment of lymphocyte expansion and adaptive immune response pathways among genes upregulated by BI-inWAX. (C) GO analysis showing enrichment of cell division pathways among downregulated genes. (D) Stacked bar plot showing relative proportions of splenic immune cell lineages. (E) Bar plots showing statistically significant differences in immune cell proportions between inWAX- and cellulose-treated mice. (F) Heatmaps and dumbbell plots showing sex chromosome-linked genes altered in both the colon and spleen. X-linked genes are in blue and Y-linked genes in red. * p < 0.05; ** p < 0.01.

To define how these transcriptional changes translate into immune cell composition, we performed immune cell fraction analysis. InWAX-treated mice showed a robust expansion of adaptive immune populations, primarily B cells and T cells (Figure 6D and 6E, Table S6). Macrophages also had considerable expansion, although to a lesser extent compared to lymphocytes. Natural killer cells were also expanded in inWAX-treated mice, but their proportion is very small to the total cells in the spleen. In contrast, dendritic cells were reduced in inWAX-treated mouse spleen, suggesting dampened activation of pro-inflammatory innate immune circuits.

Surprisingly, three Y chromosome-linked genes (*Uty*, *Kdm5d,* and *Eif2s3y,*) that were upregulated in the colon of inWAX-treated male mice were also induced in their spleens (Figure 6F). On the contrary, their X chromosome paralogs (*Utx, Kdm5c,* and *Eif2s3x*) were modestly downregulated in these mice, resulting in a marked increase in the Y-to-X expression ratio (Figure 6F). At the total gene level, inWAX increased the expression of *Eif2s3* and *Uba1* in the colon, although this did not reach statistical significance due to variability among inWAX-treated mice (Figure S3A). In the spleen, inWAX caused a significant increase in the expression level of *Uty,* but the total levels of *Utx* and *Uty* exhibited a modest decrease due to the reduced expression of *Utx*. inWAX-induced *Uba1y* only occurred in the colon (Figure 6F). Interestingly, in inWAX-treated female mice without DSS treatment, inWAX did not affect the expression of these genes in the colon (Figure S3B) This interesting finding suggest a potential sex-dependent component of BI-inWAX-mediated immune protection at both local and systemic levels.

Additionally, 12 genes induced in the spleen were also upregulated in the colon of inWAX-treated mice (Figure S3C). These shared genes included seven metabolic enzymes and regulators, the fructose transporter *Slc2a5*, the stem cell-associated factor *Piwil4*, and immune-related genes (*Apol7c* and *Retn*).

Together, these findings indicate that metabolites derived from BI-mediated inWAX fermentation reprogram immune and metabolic pathways to counteract DSS-induced inflammatory stress. This protective effect involves coordinated activation of conserved transcriptional programs across both local and systemic compartments, including modulation of sex chromosome-linked gene expression.

### InWAX fermentation ameliorates HFD-induced glucose intolerance

Given that BI-mediated inWAX fermentation increased systemic levels of anti-inflammatory and antioxidant phenolic compounds and anti-diabetic bile acids, we next examined whether these beneficial effects could be preserved in the context of HFD-induced gut dysbiosis to ameliorate metabolic dysfunction in obesity. We found BI is not a native gut resident of C57BL/6J mice, which makes this mouse strain a unique model for the study (Figure S4A). As outlined in Fig. 7A, conventionally raised male mice were first preconditioned on HFD for 8-week to induce obesity and then randomly assigned to receive HFD alone (HFD), HFD supplemented with 10% cellulose (HFD+Cellu), or HFD supplemented with 10% inWAX (HFD+inWAX) (See Material and Methods, Table S1). Cellulose served as a non-fermentable fiber control to isolate inWAX-specific effects and to control for the modest reduction in calorie density introduced by fiber inclusion (∼4% less than standard HFD).

**Figure 7.**
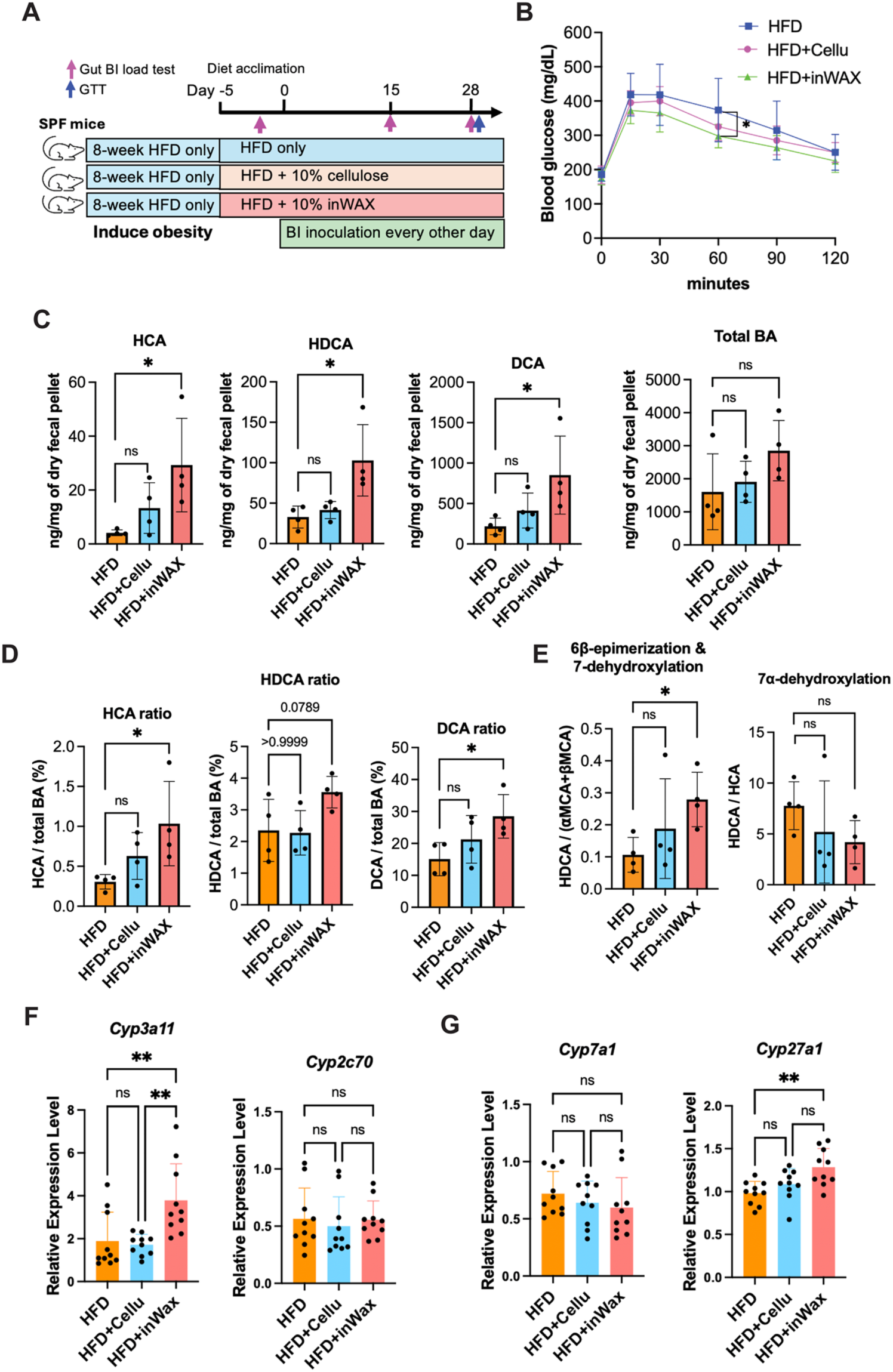
BI and inWAX improves glucose tolerance and bile acid profiles in HFD-induced obesity. (A) Schematic of the experimental workflow illustrating BI- and inWAX-based interventions in HFD-induced obese mice. All mice were inoculated with BI. (B) Glucose tolerance test results showing improved glucose clearance in inWAX-treated mice relative to HFD controls. (C) Comparison of concentrations of HCA, HDCA, DCA, and the total bile acids. (D) Comparison of the ratios of HCA, HDCA, and DCA. (E) Comparison of the ratios of HDCA to ⍺MCA and βMCA as well as HDCA to HCA. (F-G) qPCR analysis of hepatic gene expression, including *Cyp3a11*, *Cyp2c70*, *Cyp7a1*, and *Cyp27a1*. Gene expression levels were normalized to *36B4*. Each symbol represents an individual mouse and bars indicate mean values. * p < 0.05; ** p < 0.01; ns, not significant.

After a 5-day dietary acclimation period, mice in all three groups were orally inoculated with BI every other day for four weeks. BI was administered to the HFD and cellulose groups to control for BI-induced effects independent of fermenting inWAX. Body weight was monitored every other day during the first 12 days of treatment. While there is no significant difference in the weight gain between HFD+inWAX and HFD+Cellu mice, mice in these two groups consistently gained less weight than HFD-only control (Figure S4B), likely reflecting reduced caloric intake associated with fiber supplementation.

Glucose tolerance was assessed on day 29 of the treatment. HFD+inWAX mice displayed a consistent lower blood glucose level at 30-, 60-, and 90-minutes following glucose administration when compared with both HFD-only and HFD+Cellu groups, with the difference between HFD and HFD+inWAX groups at 60 minutes reaching statistical significance (Figure 7C). While cellulose-treated mice exhibited modest reductions in glucose levels relative to HFD controls, they are not statistically significant, suggesting a limited, nonspecific fiber effect. These results indicate an improvement in glucose tolerance delivered by the synergistic function of BI and WAX.

### InWAX fermentation induced health-promoting bile acid species in obese mice

To define the metabolic basis underlying the health benefits conferred by inWAX fermentation in obese mice, we examined its effects on the bile acid pool. Total 26 bile acids were evaluated by targeted quantitative assay (Table S7). InWAX-treated mice exhibited a significant increase in the concentrations of three 6-hydroxylated bile acids, HCA, hyodeoxycholic acid (HDCA), ω-muricholic acid (ωMCA), as well as deoxycholic acid (DCA) (Figure 7C). At the level of total bile acids, both inWAX- and cellulose-treated mice showed modest increases that did not reach statistical significance (Figure 7C). To determine whether the increases in HCA, HDCA, ωMCA, and DCA reflected selective enrichment rather than a general rise in total bile acids, their relative proportions within the total bile acid pool were compared across groups. Except for ωMCA, the proportions of HCA, HDCA, and DCA were substantially elevated, although the increase in HDCA narrowly missed statistical significance (Figure 7D and S5A, Table S8). Notably, both HCA and HDCA are reduced in patients with obesity, diabetes, and non-alcoholic fatty liver disease (NAFLD), and have been reported to improve glucose and lipid homeostasis and to protect against atherosclerosis ^31,76–81^. The selective expansion of these bile acid species highlights the therapeutic potential of the BI-inWAX synbiotic in ameliorating HFD-induced metabolic dysfunction by restoring health-promoting bile acid signaling.

In mice, HDCA is considered as a secondary bile acid generated through microbial biotransformation. HDCA can be produced via microbial 7-dehydroxylation and 6β-epimerization of ⍺MCA and βMCA as well as 7-dehydroxylation of HCA.^82,83^ To assess whether increased HDCA resulted from enhanced conversion of αMCA and βMCA, we compared the ratios of HDCA to these precursor bile acids. InWAX-fed mice exhibited a significant increase in the HDCA to αMCA and βMCA ratio compared with mice fed HFD alone, whereas this increase was not observed in cellulose-treated mice (Figure 7E). In contrast, the HDCA-to-HCA ratio was not increased in inWAX-treated mice. This finding suggests that HDCA production is presumably increased through enhanced microbial 7-dehydroxylation and 6-hydroxy epimerization of αMCA and βMCA.

Unlike HDCA, the biosynthetic origin of HCA in mice remains incompletely defined. Studies using human liver microsomes indicate that HCA can be produced through 6-hydroxylation of CDCA by a hepatic cytochrome P450 enzyme, CYP3A4^84,85^. Notably, HCA was upregulated by inWAX but not cellulose in BI mono-associated mice (Figure 2D), suggesting that metabolites derived from BI-mediated inWAX fermentation directly stimulated hepatic HCA production. This prompted us to examine whether BI and inWAX influence the activation of *Cyp3a11*, the murine homolog of human *Cyp3a4*. Indeed, we detected a significant increase in *Cyp3a11* expression in the livers of inWAX-fed mice. To assess whether this effect reflected broader activation of 6-hydroxylases, we also evaluated *Cyp2c70*, a hepatic gene that encodes another murine enzyme that catalyzes 6-hydroxylation of CDCA and UDCA to generate αMCA and βMCA^86^. *Cyp2c70* expression was unchanged in inWAX-treated mice (Figure 7F), which is consistent with unaltered levels of αMCA and βMCA in these mice (Figure S5B). These findings suggest that BI and inWAX selectively induce *Cyp3a11*, which presumably enhanced 6-hydroxylation of CDCA to produce HCA.

We next evaluated whether BI and inWAX influence broad hepatic bile acid synthesis pathways. Expression of *Cyp7a1*, which encodes a rate-limiting enzyme in the classical bile acid synthesis pathway, was unchanged in inWAX-fed mice (Figure 7G). *Cyp27a1*, which encodes the rate-limiting enzyme in the alternative pathway, showed a very modest increase.

Finally, we assessed bile acid deconjugation. Neither inWAX nor cellulose led to a statistically significant change in deconjugation activity when compared to HFD alone (Figure S5C).

Collectively, these findings suggest that BI and inWAX are capable of reprogramming hepatic and microbial bile acid metabolism under HFD conditions to enhance the production of health-promoting bile acids.

### InWAX fermentation induced the fecal metabolome shift

To determine how the synergy of BI and inWAX impacts gut microbial metabolic function, we used untargeted fecal metabolomics analysis to unbiasedly examine the fecal metabolome, a functional readout of microbial metabolic activity. We observed a marked remodeling of the fecal metabolic landscape in inWAX-treated mice compared with both HFD and cellulose-treated controls (Figure 8A). In total, 123 known metabolites were uniquely altered by inWAX, with 85 increased and 38 decreased (Fig. 8B;Table S9), defining a distinct microbial metabolic signature induced by BI and inWAX under HFD conditions. Structurally, the metabolites exclusively upregulated by inWAX were highly enriched for amino acids and peptides (47%) (Figure 8C, Table S10). Strikingly, ∼33% of them belonged to neurotransmitters and their precursors (Figure 8D; Table S9), suggesting that the synbiotic effect of BI and inWAX may shift protein metabolism toward enhanced neuroactive compound production and potentially strengthen the gut-brain metabolic crosstalk. The second most enriched class comprised fatty acid derivatives, predominantly oxygenated fatty acids (oxylipins) derived from long-chain polyunsaturated fatty acids (Fig. 8B;Table S9), consistent with the role of this synbiotic in rewiring lipid metabolic pathways.

**Figure 8.**
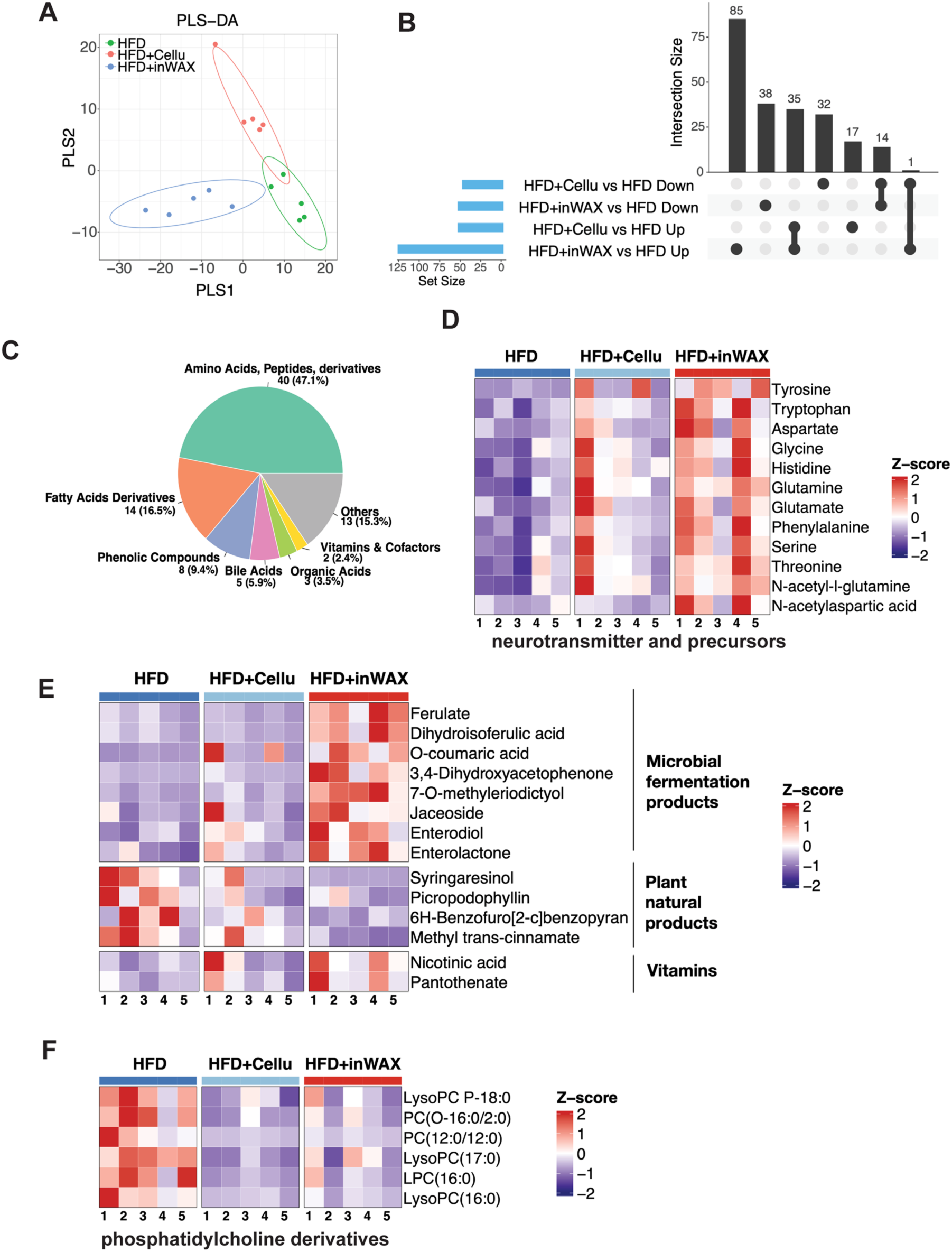
BI and inWAX rewire gut metabolic outputs under HFD conditions. (A) PLS-DA showing BI-inWAX induced a metabolite profile distinct from cellulose. (B) The UpSet plot showing metabolites uniquely or commonly altered among the indicated groups. (C) Pie chart showing structural classification of metabolites uniquely induced by BI-inWAX. (D) Heatmap showing increased production of neurotransmitters and their precursors. (E) Heatmap showing BI-inWAX-induced increases in microbial fermentation-derived bioactive phenolic compounds and vitamins, alongside reduced levels of plant-derived polyphenols. (F) Heatmap showing BI-inWAX downregulated phosphatidylcholine derivatives.

In line with BI-mediated liberation of ferulic acid from inWAX, we observed a robust increase in ferulate, a salt form of ferulic acid, and its derivative dihydroisoferulic acid (Figure 8E). Notably, 6 other phenolic compounds with well-documented anti-inflammatory and antioxidant activities^87–89^, were also significantly elevated. This includes 2 simple phenolics (o-coumaric acid and 3,4-dihydroxyacetophenone), 2 flavonoids (7-O-methyleriodictyol and jaceoside), and 2 enterolignans (enterolactone and enterodiol). While ferulic acid and o-coumaric acid are typically ester-linked to the arabinoxylan backbone and are therefore likely direct products of inWAX fermentation, the remaining phenolic compounds represent microbial biotransformation products of dietary polyphenols. Concomitantly, multiple complex plant polyphenols were significantly reduced, among which syringaresinol can be microbially converted to enterolactone and enterodiol^90^. These reciprocal changes indicate that the BI–inWAX synbiotic not only releases health-promoting metabolites through inWAX fermentation (e.g., ferulic acid) but also enhances microbial conversion of dietary polyphenols into bioactive metabolites. Notably, this metabolic rewiring was observed exclusively in BI-inWAX treated mice, supporting a model wherein BI-inWAX promotes cooperative, cross-feeding-dependent gut community function.

Additionally, inWAX fermentation significantly elevated nicotinic acid (vitamin B3) and pantothenate (vitamin B5) (Figure 8E), two essential cofactors for cellular energy metabolism. While nicotinic acid can be synthesized by both host and microbes, pantothenate is exclusively produced by gut microbes, further supporting a microbiota-dependent reprogramming of metabolic capacity.

Unlike upregulated metabolites, the metabolites exclusively downregulated in InWAX-treated mice were structurally heterogeneous (Table S9). Besides the four aforementioned plant polyphenols, it includes sphingolipids and glycerophospholipids, suggesting a role of BI and inWAX in modulating lipid metabolism.

We also identified 49 metabolites altered in both inWAX- and cellulose-treated mice (35 increased, 14 decreased), likely reflecting shared fiber-dependent effects. Most of the co-upregulated metabolites are amino acids and short peptides (Table S9), indicating that both fermentable and nonfermentable dietary fiber enhanced protein metabolism. Interestingly, the co-downregulated metabolites include 6 phosphatidylcholine derivatives involved in membrane composition regulation and lipid mediator signaling pathways (Fig 8F). Notably, 3 of them, LysoPC(17:0), LPC(16:0), and LysoPC(16:0), are also downregulated in the serum of inWAX-treated BI mono-associated mice (Figure 2G), suggesting that BI-mediated inWAX fermentation has a direct role in reducing their production, and this function is preserved under the gut microbiota of HFD-induced obesity.

Additionally, 49 metabolites were uniquely altered by cellulose, with 17 increased and 32 decreased, indicating that fermentable inWAX and non-fermentable cellulose exert distinct metabolic effects under high-fat diet conditions. 8 of the 17 cellulose-upregulated metabolites were short peptides, suggesting that, despite being non-fermentable, cellulose may still influence protein metabolism, albeit through pathways distinct from those activated by fermentable inWAX. Except for a single sulfated dietary polyphenol metabolite, 3-(3-hydroxyphenyl)propionic acid sulfate, cellulose did not increase the level of beneficial phenolic compounds and bile acids as inWAX.

Together, these findings indicate that the synergistic interaction between BI and inWAX is critical for conferring health-promoting effects under HFD-induced obesity. This synergy enhances microbial production of anti-inflammatory and antioxidant phenolic metabolites. It also promotes the selective expansion of anti-diabetic and anti-steatotic bile acids, HCA and HDCA, through remodeling hepatic and microbial bile acid transformation pathways. These findings highlight the potential of leveraging this synbiotic to reprogram host metabolism and immunity for preventing and mitigating metabolic and inflammatory diseases.

## Discussion

The modern low-fiber diets have driven the loss or depletion of fiber-fermenting microbes, thereby attenuating the health benefits of fiber consumption. Synergistic synbiotics, which pair defined microbial strains with their preferred fiber substrates, offer a promising strategy to restore these functions. However, the rational design of such interventions is constrained by insufficient understanding of microbial fiber-degrading capacities and the host-relevant bioactivities of fermentation-derived metabolites.

In this study, we present compelling evidence that BI is a key microbial mediator of dietary fiber-driven metabolic, immune, and potentially neuronal benefits. We show that the synergistic interaction between BI and its dietary fiber substrate, inWAX, selectively enhances the production of anti-diabetic and anti-steatotic bile acid species, as well as anti-inflammatory and antioxidant phenolic compounds even in the context of HFD-induced gut microbiota. Further, it remodels protein metabolism toward enhanced neuroactive compound production, which potentially strengthens the gut-brain metabolic crosstalk. These microbial metabolites are accompanied by coordinated transcriptional reprogramming in the colon and spleen that improves circadian rhythm regulation, lipid metabolism, and immune defense.

A particularly notable finding is the selective expansion of health-promoting bile acids, HCA, HDCA, and TUDCA. These bile acids have been shown to ameliorate metabolic disorders such as diabetes, atherosclerosis, and non-alcoholic fatty liver disease (NAFLD) ^31,77–79^. Reduced systemic levels of HCA and HDCA are found in individuals with obesity and diabetes, further underscoring their therapeutic relevance ^31,76^ TUDCA, which is clinically approved for primary biliary cholangitis due to its cytoprotective properties, has also shown promise in obesity, IBD, and neurodegenerative diseases ^29,30,91–95^. The selective expansion of these beneficial bile acids by the synergistic effects of BI and inWAX, even under HFD-induced obesity, highlights the translational potential of this synbiotic strategy for metabolic disease intervention.

Mechanistically, how BI-mediated inWAX fermentation enhances the production of these bile acid species is of great interest. In BI mono-associated mice, inWAX increased systemic TUDCA and HCA levels compared with cellulose, suggesting that BI modifies hepatic bile acid metabolism through fermenting inWAX, likely by releasing metabolites that reprogram hepatic bile acid biosynthetic pathways. In line with this idea, we observed an upregulation of *Cyp3a11*, a gene encodes a 6-hydroxylase that converts CDCA to HCA in human liver microsomes. The specific metabolite(s) responsible for inducing *Cyp3a11* expression warrants further investigation.

In contrast, the increase of HDCA by BI and inWAX occurs only in the presence of the complex gut microbiota, suggesting that its biotransformation needs the cooperative activity of additional gut microbial species. In mice, HDCA can be produced by microbial 7-dehydroxylation and 6-hydroxy epimerization of ⍺MCA and βMCA. The significantly increased ratio of HDCA to ⍺MCA and βMCA in inWAX-treated mice support a model system wherein BI-inWAX provide nutrients that cross-feed gut microbes that convert ⍺MCA and βMCA to HDCA. In line with this idea, a study in gnotobiotic rats found that a Gram-positive microbe termed HDCA-1, is capable of converting ⍺MCA and βMCA to HDCA via 7-dehydroxylation and 6-hydroxy epimerization only in the presence of a growth factor-producing strain.^83^ A more recent study in mice shows that *Bacteroides ovatus*, a member of *Bacteroides* spp., promote HDCA production by supporting the growth of 7-dehydroxylating bacterium *Clostridium scindens*.^96^ Future experiments that co-culture BI and HDCA-producing microbes in the presence of inWAX could help address this model hypothesis.

In addition to selectively expanding beneficial bile acid species in HFD-induced obese mice, BI and inWAX also enhance the microbial biotransformation of dietary plant polyphenols into a spectrum of bioactive metabolites, including simple phenolics, flavonoids, and enterolignans. These compounds are well documented for their potent anti-inflammatory and antioxidant activities^87–89^. This enhanced biotransformation suggests that BI and inWAX not only generate specific metabolites such as ferulic acid with direct therapeutic potential, but also broadly augment gut microbiota-mediated metabolic processes. As a result, this synbiotic intervention substantially amplifies the host’s capacity to extract and utilize the health-promoting benefits of dietary fiber.

An unexpected and conceptually important finding is the induction of core circadian clock genes in the intestine following BI-mediated inWAX fermentation. The intestinal circadian clock is critical for mucosal barrier integrity, immune responses, metabolism, and gut microbiota function. Disrupted circadian rhythm has been associated with a higher risk and more aggressive course of IBD ^97,98^. Reduced colonic expression of *Per1* and *Per2* has been observed in patients with IBD ^99,100^, and mice lacking these genes exhibit impaired antimicrobial secretion and heightened sensitivity to DSS-induced colitis ^101^. Our findings uncover a previously unrecognized link between microbial fiber fermentation and intestinal circadian regulation, potentially connecting microbial metabolism with gut–brain–immune axis signaling.

Another interesting finding from this study is BI-mediated inWAX fermentation induces robust activation of Y chromosome-linked genes *Eif2s3y, Kdm5d,* and *Uty,* in both the colon and spleen during immune defense against DSS-induced colitis. These genes function in RNA translation and epigenetic regulation, processes essential for stress and immune responses. Their induction suggests a potential sex-biased component in benefiting from fiber fermentation.

Together, our findings redefine the prevailing paradigm of fiber-microbiota-host interactions by establishing that targeted fiber fermentation drives substantial health benefits through reprogramming hepatic and microbial bile acid metabolism, enhancing the production of bioactive phenolic compounds, and activating protective transcriptional programs in the colon, spleen, and liver. Our work, thus, establishes a rational framework for the design of the BI-inWAX synbiotic to prevent and treat inflammatory and metabolic diseases.

## Acknowledgments

We thank Dr. Bo Wang, Ms. Tiantian Yang, and Mr. Zhiming Zhao for technical assistance with metabolic parameter measurements, Dr. Kun Wang for suggestions on immune cell fraction analysis, and Dr. Nicole Koropatkin (University of Michigan) for advice on testing BI function. We thank Dr. Alexander Ulanov, Dr. Michael Frano, and Ms. Wei Lu at the Metabolomics Core of the Roy J. Carver Biotechnology Center, UIUC, for metabolomics analysis. We thank the DNA sequence service core at the Roy J. Carver Biotechnology Center for RNA-sequencing. We thank Karen Doty at the Histology Laboratory for assistance with histological staining.

---

**Figure S1.**
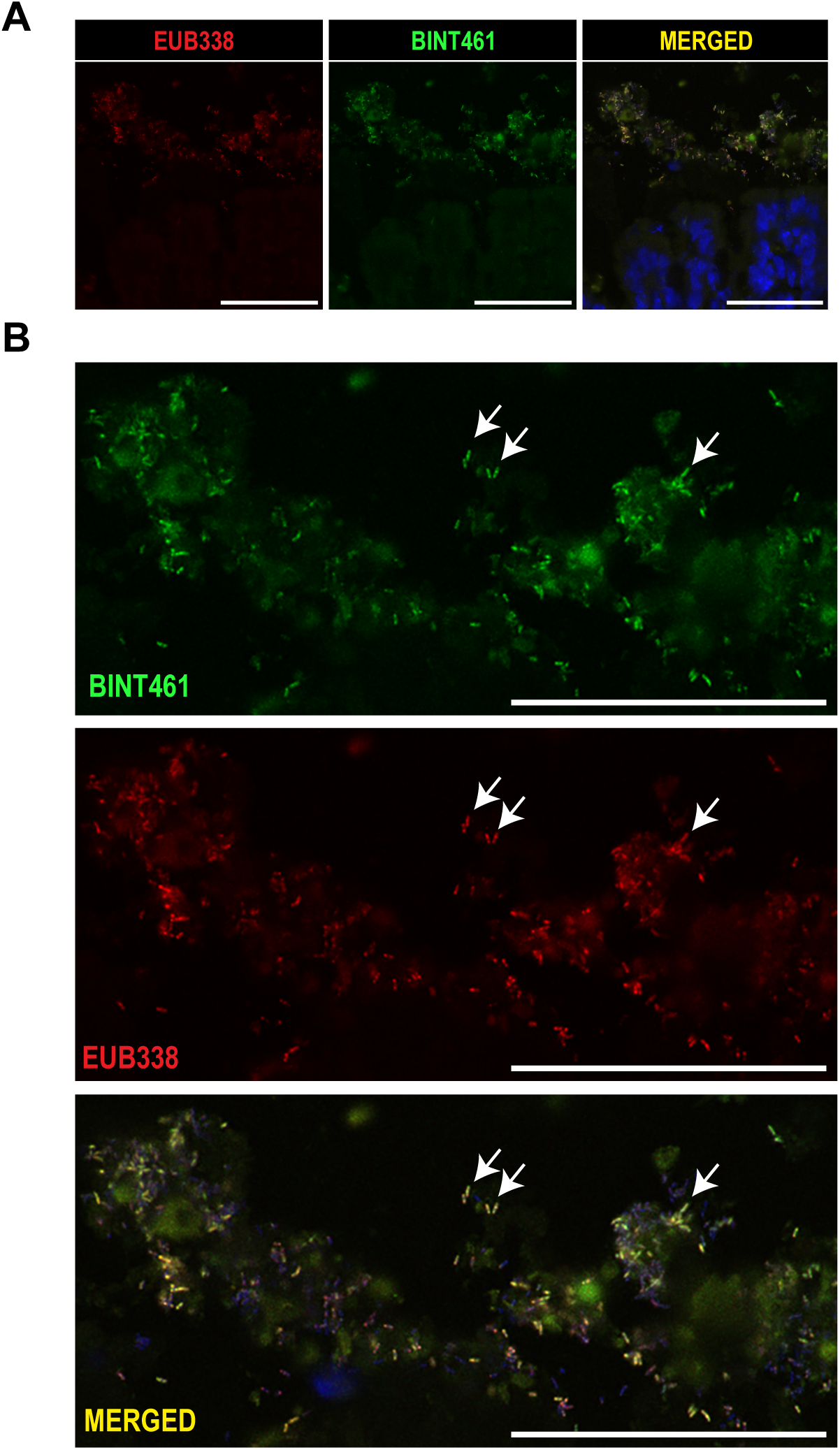
FISH detection of BI in the colonic lumen. (A) BI was detected using the BI-specific probe BNT461. The universal bacterial probe EUB338 served as a positive control. DAPI staining (blue) in the merged image labels colonic epithelial cell nuclei. (B) Higher-magnification views corresponding to the regions shown in (A). Arrows point to some BI. Scale bars, 50 µm.

**Figure S2.**
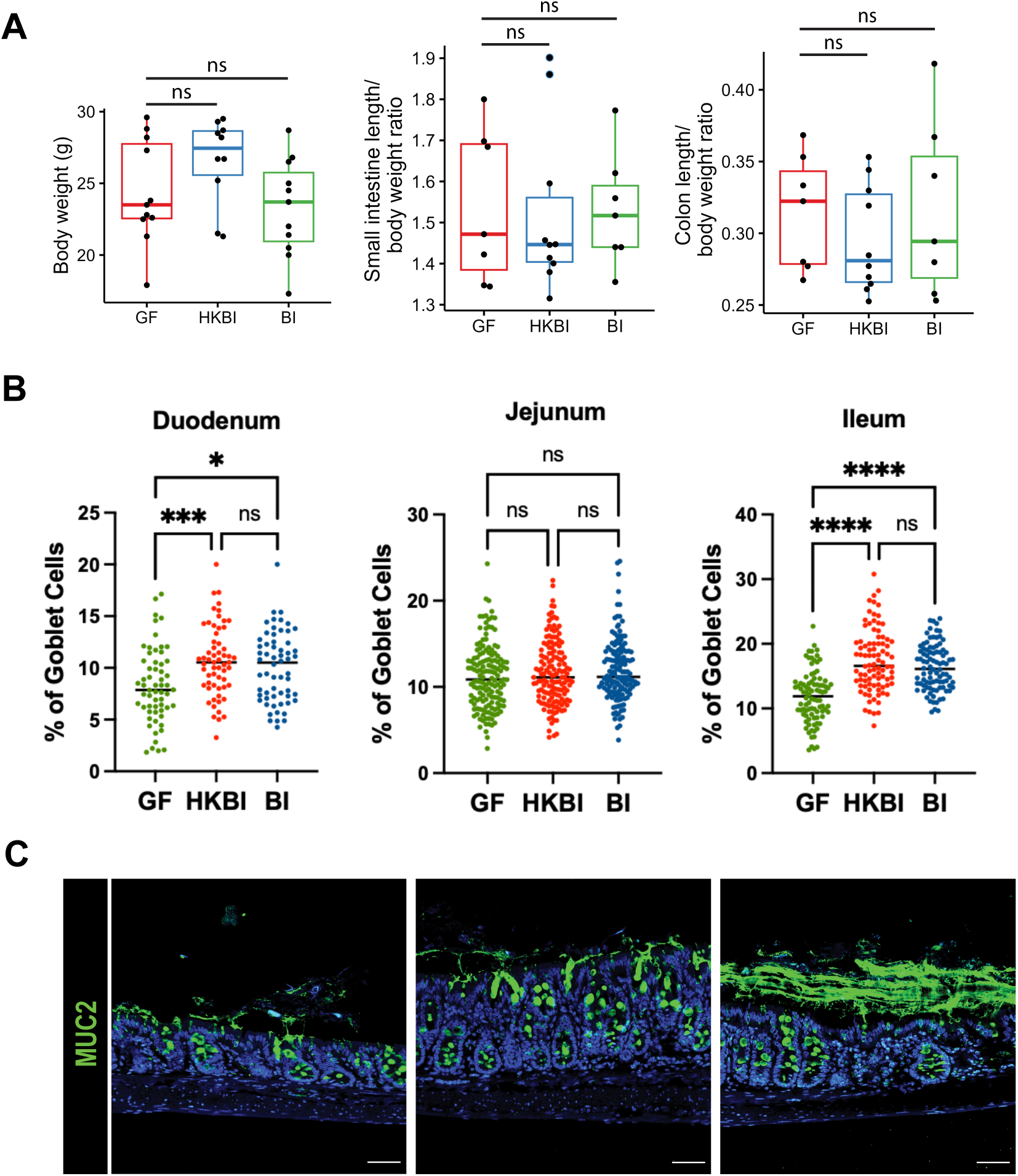
Physiological impact of BI monoassociation. (A) Quantification of body weight and the ratio of small intestine and colon length to body weight. (B) Quantification of goblet cells as a percentage of total colonic epithelial cells. (C) Representative immunofluorescence staining of MUC2 in the colon. Scale bars, 50 μm. ** p < 0.01; *** p < 0.001; **** p < 0.0001. ns, not significant.

**Figure S3.**
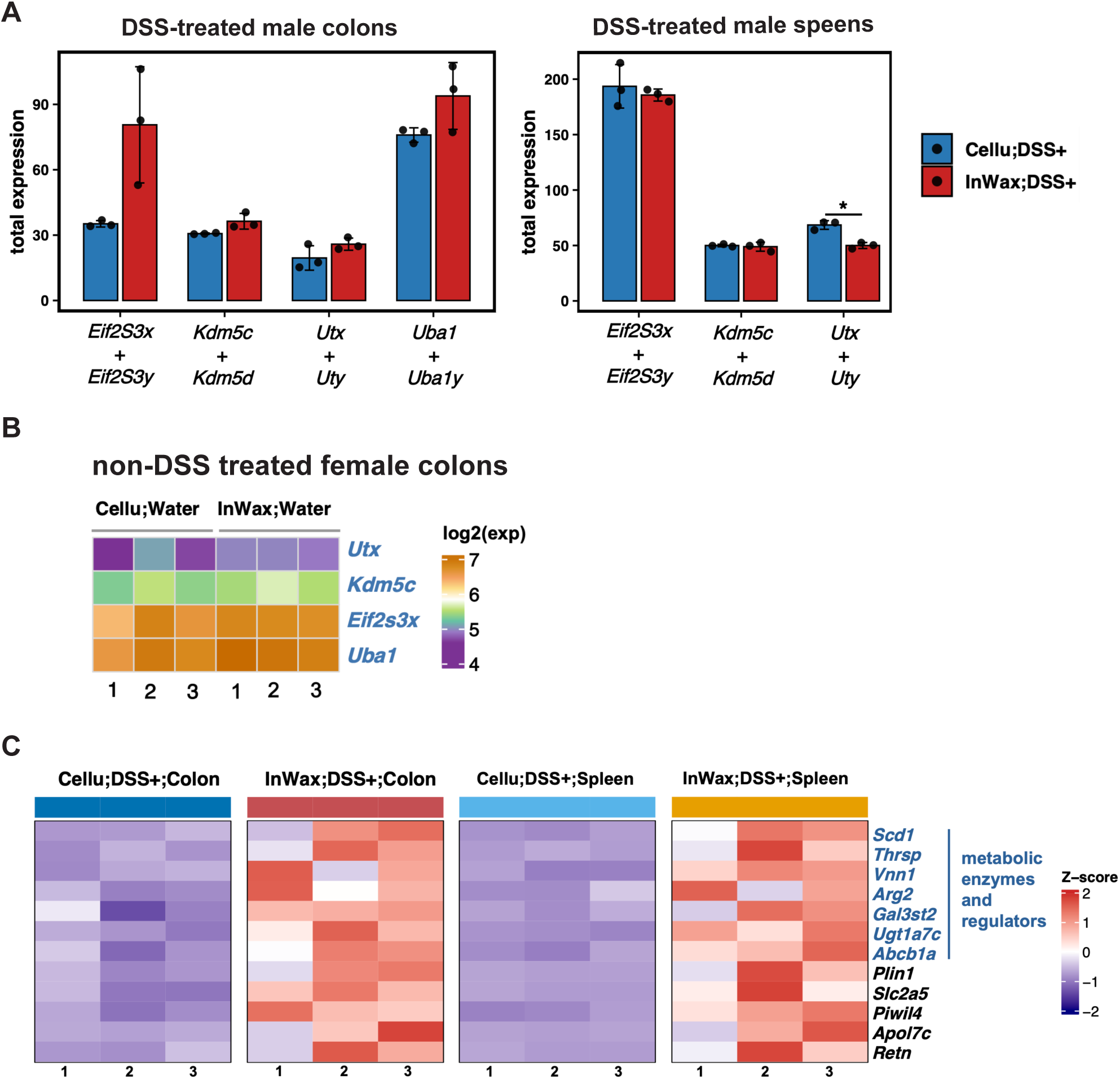
BI and inWAX induced transcriptomic changes in the colon and spleen. (A) Comparison of total expression levels of sex chromosome-linked paralog genes in the colons and spleens in response to DSS. (B) Heatmap showing unaltered X chromosome-linked genes between the colons of female mice treated with cellulose and inWAX. (C) Heatmap showing metabolic genes upregulated by inWAX in the colon and spleens in response to DSS treatment. * p < 0.05. ns, not significant

**Figure S4.**
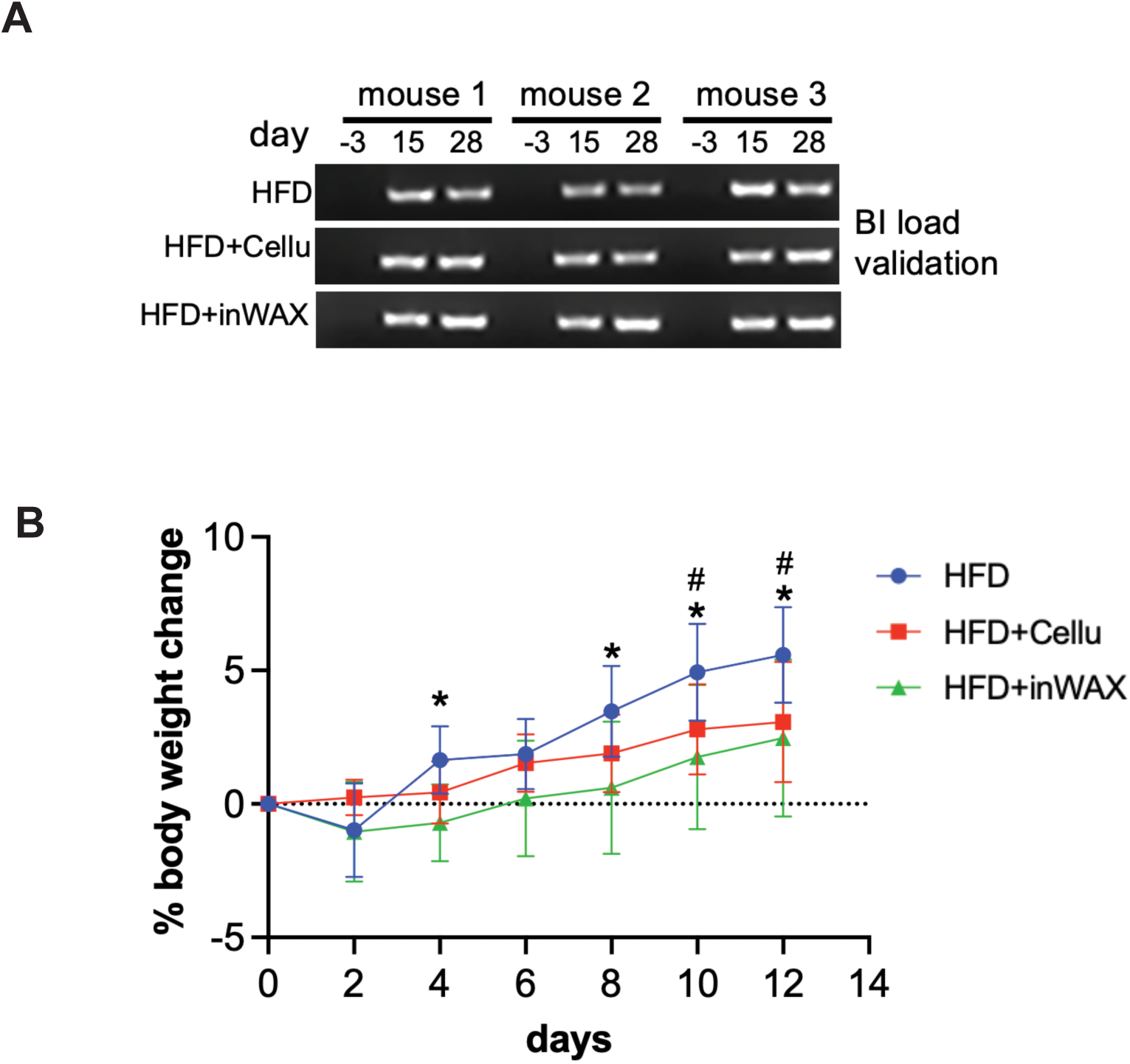
BI abudnace validation and difference in mouse body weight changes. (A) PCR validation of BI abundance in fecal samples of 3 mice in each group at the indicated time points. The first day of BI inoculation was designated as day 0. BI was undetectable prior to inoculation (day −3), indicating it is not a native resident of B6 mice.(B) Comparison of body weight change (%) during the first 12 days of BI inoculation. * indicates significance between HFD+inWAX and HFD. # indicates significance between HFD+cellu and HFD. * p < 0.05; # p < 0.05.

**Figure S5.**
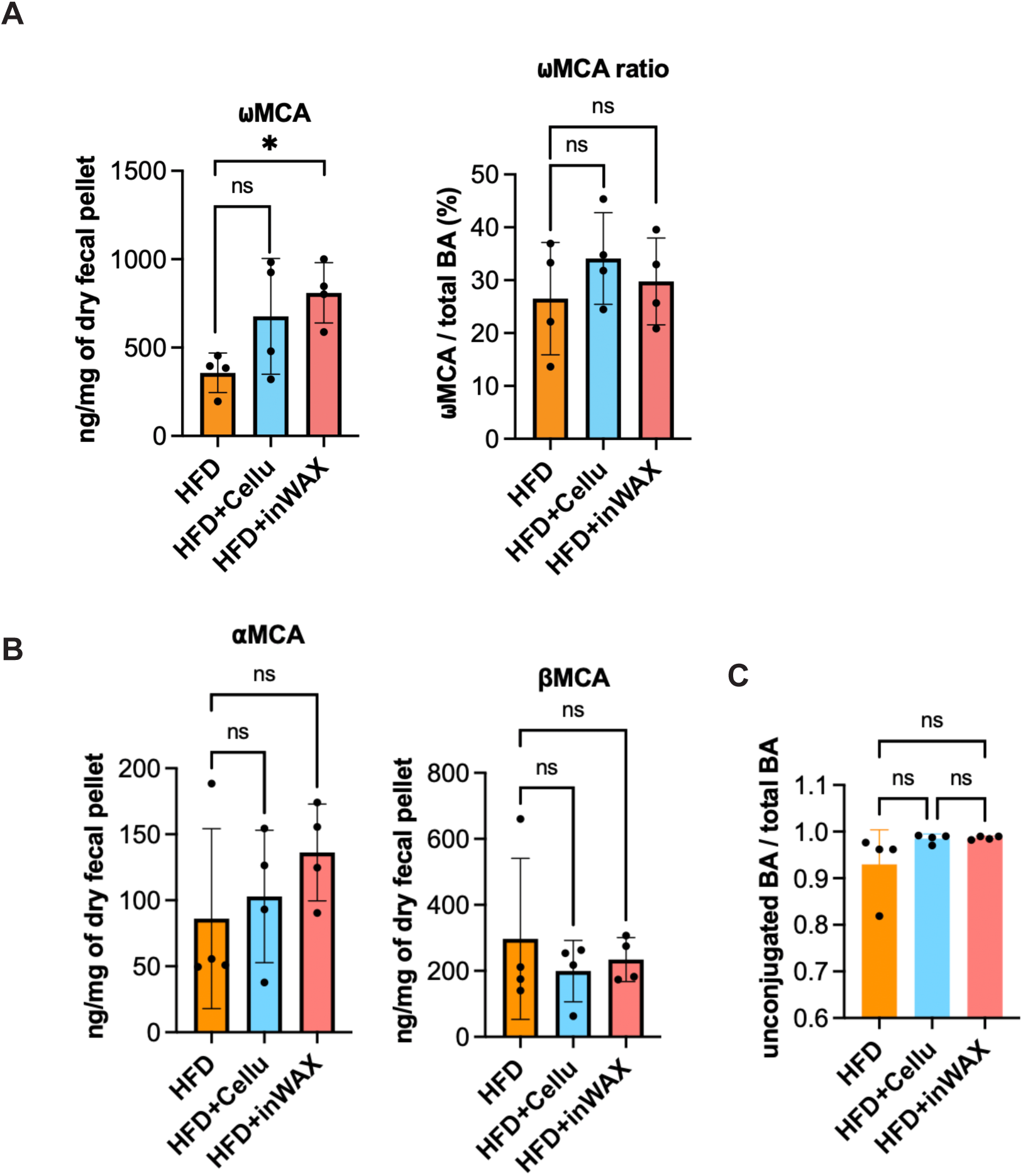
BI and inWAX induced changes in fecal bil acid profile. (A) Comparison of ωMCA concentrations (ng/ mg) and ωMCA proportions. (%). (B) Comparison of aMCA and bMCA concentrations (ng/ mg). (C) Comparison of the ratios of unconjugated bile acid (BA) to total BA. * p < 0.05. ns, not significant

